# Targeted Delivery of Chloroquine to Plasmacytoid Dendritic Cells Enhances Inhibition of the Type I Interferon Response

**DOI:** 10.1101/2021.06.09.447773

**Authors:** Marilyn E. Allen, Amit Golding, Violeta Rus, Nicholas B. Karabin, Sophia Li, Chamille J. Lescott, Sharan Bobbala, Evan A. Scott, Gregory L. Szeto

## Abstract

Systemic lupus erythematosus (SLE) causes damaging inflammation in multiple organs via the accumulation of immune complexes. These complexes activate plasmacytoid DCs (pDCs) via TLR7 and TLR9, contributing to disease pathogenesis by driving secretion of inflammatory type I IFNs. Antimalarial drugs, such as chloroquine (CQ), are TLR antagonists used to alleviate inflammation in SLE. However, they require ~3 months of continuous use before achieving therapeutic efficacy and can accumulate in the retinal pigment epithelium with chronic use resulting in retinopathy. We hypothesized that poly(ethylene glycol)-*b*-poly(propylene sulfide) (PEG-*b*-PPS) filamentous nanocarriers, filomicelles (FMs) could improve drug activity and reduce toxicity by directly delivering CQ to pDCs via passive, morphology-based targeting. Healthy human PBMCs were treated with soluble CQ or CQ-loaded FMs, stimulated with TLR agonists or SLE patient sera, and type I IFN secretion was quantified via multi-subtype IFN-α ELISA and *MX1* gene expression using real-time RT-qPCR. Our results showed that 50 µg CQ/mg FM decreased *MX1* expression and IFN-α production after TLR activation with either synthetic nucleic acid agonists or immune complex rich sera from SLE patients. Cellular uptake and biodistribution studies showed that FMs preferentially accumulate in human pDCs *in vitro* and in tissues frequently damaged in SLE patients (*i.e.,* liver and kidneys) while sparing the eye *in vivo*. These results showed that nanocarrier morphology enables drug delivery, and CQ-FMs may be equally effective and more targeted than soluble CQ at inhibiting SLE-relevant pathways.

## Introduction

Systemic lupus erythematosus (SLE) is an immune complex-mediated autoimmune disease characterized by dysregulation of the innate and adaptive immune system resulting in loss of self-tolerance. From 2000 to 2015, SLE was among the leading causes of death among young women aged 15-24 in the United States (1). Women represent 90% of patients, leading to a strong female bias in disease demographics (2, 3). Clinical manifestations are heterogeneous with varying disease severities, organ involvement, and cellular abnormalities (4, 5). The clinical course is unpredictable, with frequent flares, which contributes to both delayed diagnosis (an estimated six years after initial presentation) (6) and difficulties in treatment. Circulating immune complexes, consisting of autoantibodies and endogenous antigen, are deposited in tissues leading to inflammation and end-stage organ damage. Lupus nephritis is one of the leading causes of morbidity and mortality in SLE, developing in up to 50% of patients (4). Plasmacytoid dendritic cells (pDCs) are activated by immune complexes sensed via toll-like receptors (TLR)-7 and -9, leading to production of interferon (IFN)-α, a major driver in SLE pathogenesis (7, 8). Pro-inflammatory IFN-α is upregulated in 50-75% of adult SLE patients (9) and can promote suppression of regulatory T cells (10), B cell differentiation to plasma cells, and the production of autoantibodies from those plasma cells (11), resulting in a positive feedback loop driving autoimmunity. Attenuating this pro-inflammatory type I IFN response is key to treating SLE.

Emerging therapeutic strategies targeting type I IFN include anifrolumab, a monoclonal antibody blocking type I IFN receptor subunit 1 (12); IFN-α kinoid, an inactivated IFN-α coupled to a carrier protein (13); and pDC inhibition via cell surface receptor blood DC antigen 2 (BDCA-2/CD303) (14). None have been approved for SLE patients, and broad type I IFN blockade may blunt antiviral immunity, resulting in an increased risk of infection and infection-related complications (15, 16). Antimalarial drugs, such as chloroquine (CQ; brand name, Aralen) and hydroxychloroquine (HCQ; brand name, Plaquenil), have been used for the treatment of SLE since the 1950s (17). They are the cheapest and most frequently prescribed first line, non-steroid disease-modifying antirheumatic drug (18). Antimalarial drugs act by binding to nucleic acids of immune complexes to mask key binding epitopes, preventing TLR7 and TLR9 activation and subsequent type I IFN responses (19–21). Antimalarial drugs are safe during pregnancy (22, 23), improve survival (24), reduce disease activity (17), and are well-tolerated with adjunctive immunomodulatory treatment (25). However, significant challenges during treatment include prolonged (> 3 months) use required before achieving therapeutic efficacy (26), poor patient compliance (27), and risk of retinopathy with chronic use (28, 29). We sought to enhance the potency of antimalarial drugs and address key challenges using targeted delivery.

In this work, we used filamentous nanocarriers composed of an oxidation-responsive block copolymer, (poly(ethylene glycol)-*b*-poly(propylene sulfide) (PEG-*b*-PPS) to target CQ to pDCs and directly inhibit type I IFN activation by TLR-driven signaling. Self-assembled PEG-*b*-PPS nanocarriers are non-immunogenic and non-inflammatory, exhibiting neither anti-PEG antibodies nor complement activation in mice and non-human primates (30, 31). The hydrophobic PPS block facilitates retention and controlled release of hydrophobic drugs such as CQ (32). Oxidation converts the hydrophobic PPS block to hydrophilic poly(sulfoxides) or poly(sulfones), leading to nanocarrier disassembly and release of drug payload (30). The hydrophilic PEG fraction controls the morphology of the self-assembled structures (32). Previous studies have identified morphology as a passive mechanism for altering cellular targeting and biodistribution. In particular, PEG-*b*-PPS filamentous worm-like micelles, or filomicelles (FMs), preferentially accumulated in splenic pDCs after subcutaneous injection (33). Passive, morphology-based targeting via FMs avoids the use of cell-specific ligands and improves blood circulation times (34). Furthermore, scalable self-assembly and loading of FMs can be successfully achieved (35, 36). We hypothesized that targeted delivery of the antimalarial drug, CQ, to pDCs via nanocarriers will enhance inhibition of type I IFN, reduce drug toxicity by focusing delivery to specific cells and tissues, and increase efficacy per dose (**Figure 1**). Our work found that CQ-loaded FMs were equivalent or more effective than soluble CQ or CQ-loaded spherical PLGA nanocarriers in decreasing *MX1* gene expression and IFN-α production by purified TLR7 and TLR9 agonists, and SLE patient sera. Combined with their preferential accumulation in pDCs and tissues of increased inflammation in SLE patients (*e.g.*, kidneys and liver) and minimal accumulation in the eyes, CQ-loaded FMs may be a novel, more effective, and more targeted formulation for treating SLE.

**Figure 1.**
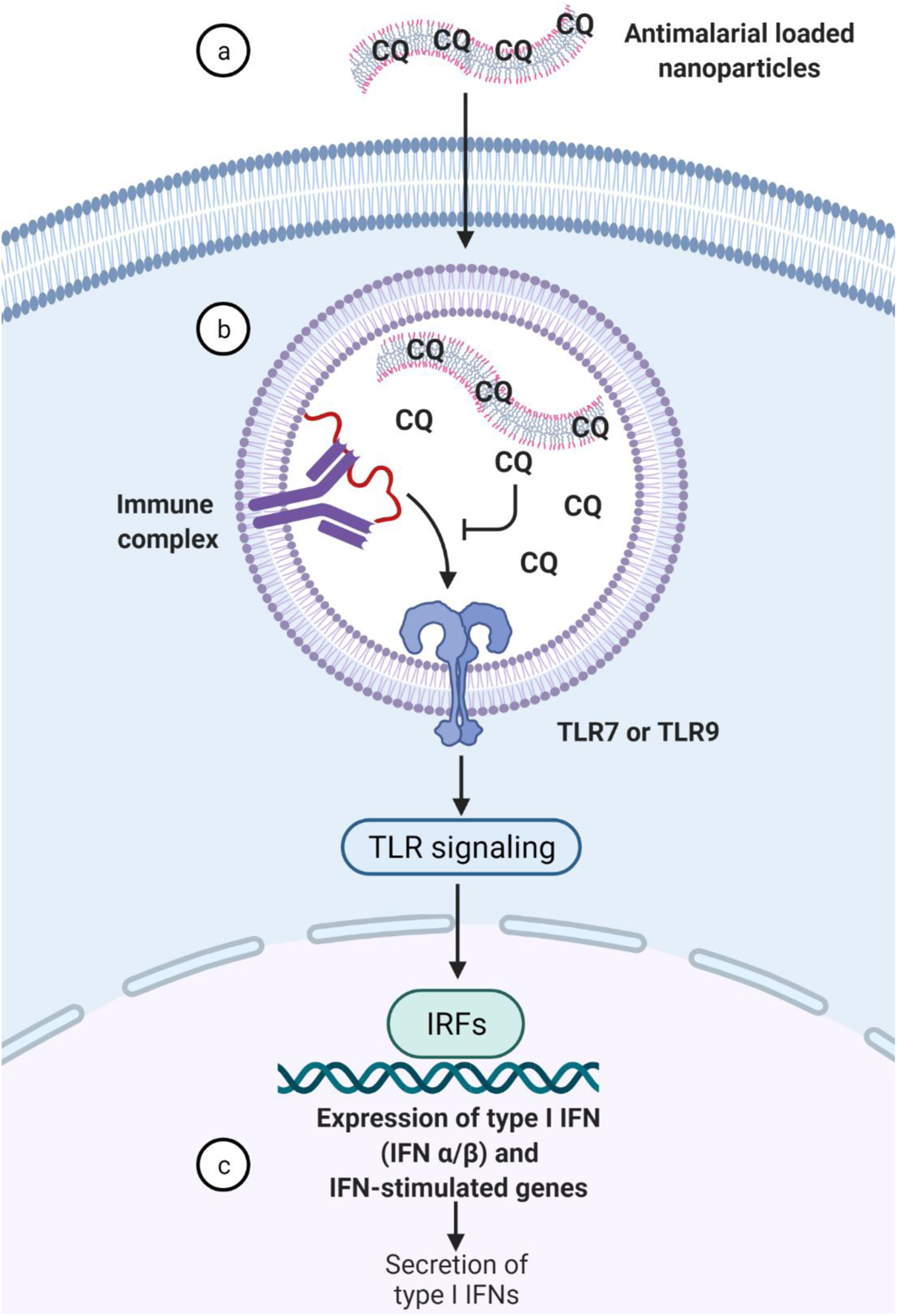
Graphical abstract. (A) Antimalarial drug-loaded polymeric nanocarriers traffic into endosomes where intracellular TLR7 and TLR9 are located. (B) Controlled release of antimalarial drug, such as chloroquine (CQ), inhibits TLR7 and TLR9 activation by masking the binding epitope of nucleic acids contained on immune complexes. (C) TLR inactivation prevents downstream signaling of type I interferon (IFN) and IFN-stimulated genes and production of pro-inflammatory cytokines, such as IFN-α, a major driver of SLE pathogenesis. Created with BioRender.com.

## Results

### Synthesis and characterization of chloroquine-loaded nanocarriers

FMs and control spherical PLGA nanocarriers were synthesized by thin-film hydration and emulsion/solvent evaporation, respectively. Nanocarrier morphology was verified via small angle x-ray scattering (SAXS). As expected, SAXS analysis of FMs best fit the scattering profile to a flexible cylinder model (**Figure S1**). The aspect ratio was calculated, and drug loading had no significant effect when comparing unloaded (χ^2^ = 0.008) versus CQ-loaded (χ^2^ = 0.0012) FMs (**Figure 2A**). Key limitations of SAXS analysis methods are the assumptions of constant shape and homogeneity in a given sample. Direct visualization by TEM was used to complement SAXS and reveal potential variations in morphology (37). Representative images showed that the morphology of unloaded and CQ-loaded FMs were consistent with micron length and ~50 nm cross-sectional diameter (**Figure 2B**). Dynamic light scattering determined control spherical PLGA nanocarriers had an average hydrodynamic diameter of 662.5 nm and 0.272 polydispersity index (PDI) for blank nanocarriers compared to 562.6 nm and 0.221 PDI for CQ-loaded nanocarriers. Surface charge is an important nanocarrier characteristic because it is a major determinant of serum protein adsorption and cellular internalization by the mononuclear phagocyte system, which consists mainly of tissue-resident macrophages (38). The zeta potential of blank and loaded nanocarriers was determined to be negative (**Figure 2C**). Neutral or anionic nanocarriers are less likely to adsorb serum proteins, be sequestered by tissue-resident macrophages, and have short serum half-lives in comparison to cationic nanocarriers (39, 40). These data confirmed that the physical (morphology, size) and biochemical (charge) properties of nanocarriers were not affected by drug loading.

**Figure 2.**
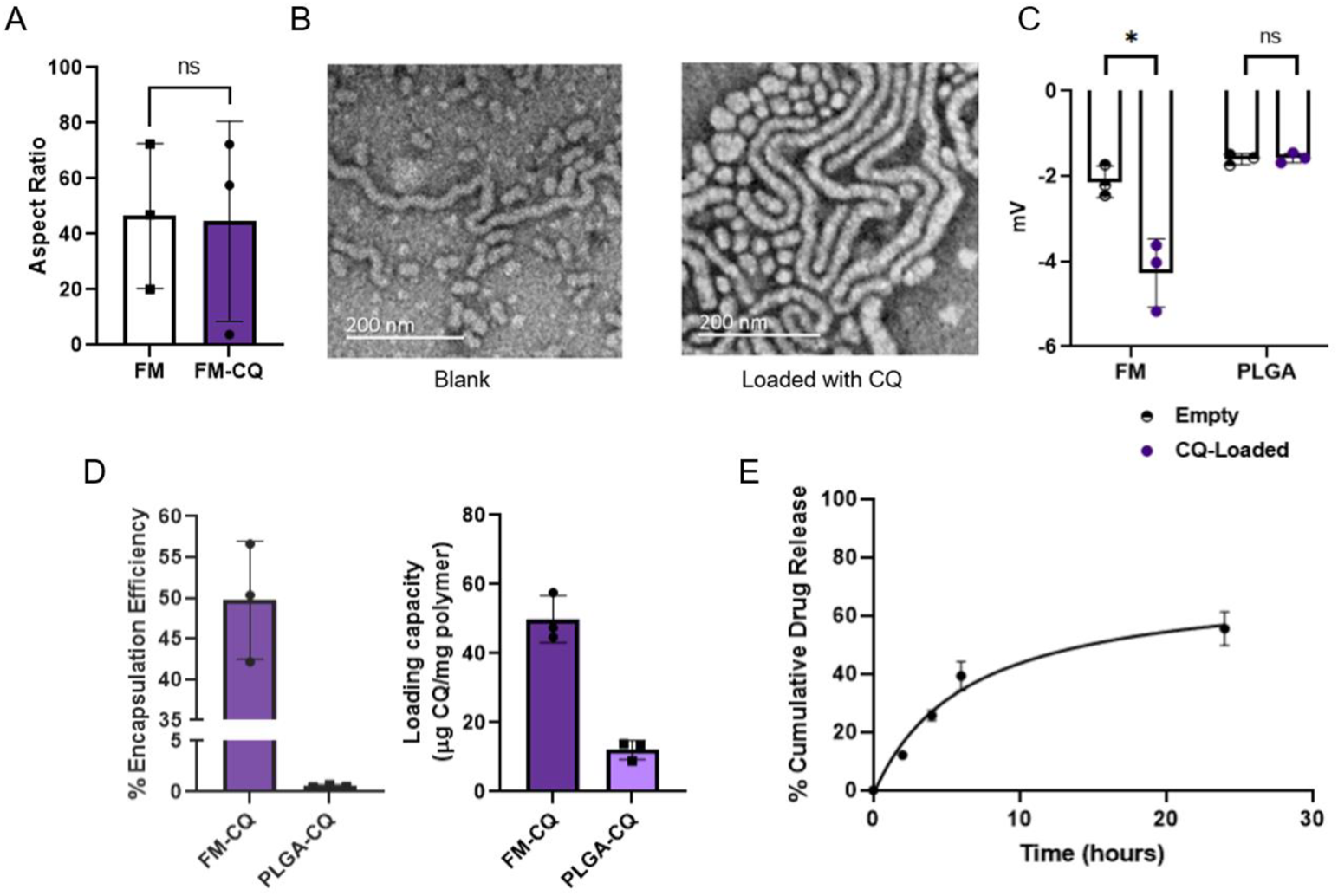
CQ loading did not alter physicochemical properties of nanocarriers. (A) Aspect ratio (width/height) of blank FMs (*n* = 3) and CQ-loaded FMs (FM-CQ) (*n* = 3). Length and diameter were measured by SAXS using the flexible cylinder model and the following parameters: 2 µm cylinder length, 150 nm persistence length, and 8 nm PPS core radius. Paired t-test for statistical analysis. (B) Representative images from TEM showed blank (left) and CQ-loaded (right) FMs at 12 mg/mL in 1X PBS. Samples were stained with 2% uranyl acetate. (C) Zeta potential (*n* = 3) of CQ loaded and blank (left) FMs or (right) PLGA. Data were shown as means with error bars representing standard deviation. Paired t-test was used for statistical analysis. Statistical significance: *p ≤ 0.05. All samples were measured in 1X PBS solution (pH 7.4) at 1 mg/mL. (D) Loading capacity (*n* = 3) and encapsulation efficiency (*n* = 3) of CQ-loaded (left) FMs or (right) spherical PLGA nanocarriers. All samples were measured in 1X PBS solution, pH 7.4 at 1 mg/mL. Data show mean and error bars represent standard deviation. (E) CQ drug release profile from FMs. CQ-loaded FMs (*n* = 3) were placed in 10K MWCO dialysis tubing in 1X PBS buffer plus 1% BSA for 0, 2, 4, 6, and 24 h at 37°C, 5% CO_2_. Each time point shows the average percentage cumulative drug release; error bars represent standard deviation.

Next, we characterized the loading and release properties of CQ. Loading capacity and encapsulation efficiency were determined by dissolving CQ-loaded nanocarriers in DMSO and quantifying CQ by UV/Vis spectrophotometry. The average loading capacity was 49.96 ± 5.529 μg CQ/mg particle (mean ± standard deviation) for FMs (**Figure 2D**) and 12.07 ± 2.255 μg CQ/mg particle for spherical PLGA nanocarriers (**Figure 2D**). The encapsulation efficiency for CQ was ~50% for FMs and ~0.6% for spherical PLGA nanocarriers (**Figure 2D**). For kinetic release studies, FMs were incubated *in vitro* for up to 24 h, and a characteristic burst release was observed followed by a plateau at ~50% release of encapsulated CQ (**Figure 2E**). Overall, these results demonstrated that CQ efficiently loaded into FMs and enabled controlled release.

### PEG-b-PPS FMs selectively target pDCs in vitro

Nanocarrier size and shape are known to significantly alter their delivery and biodistribution *in vitro* and *in vivo* (38). We evaluated whether FMs could address two key design needs: 1) targeting and preferential accumulation in pDCs *in vitro* and 2) biodistribution favoring major target organs in SLE while avoiding off-target effects in the eye *in vivo*. FMs were loaded with the lipophilic fluorescent dye DiD for tracking and added to cultures of human PBMCs *in vitro.* Flow cytometry was used to determine cellular targeting and association after 6, 24, and 48 h. pDCs represent an average of 0.29% of all cells in healthy human PBMCs but accumulated more FMs than B and T cells after 48 h (**Figure 3A**). DiD+ cells were quantified, and their median fluorescent intensity calculated to estimate both the fraction of cells taking up nanocarriers as well as the amount of nanocarriers internalized per cell. pDCs were consistently associated with nanocarriers compared to more abundant cells (12.6% DiD+ of T cells) and total live cells (98.08% of PBMCs) (**Figure 3A**). The intensity of DiD in pDCs was also highest among all analyzed cell types (average MFI of 25.73 after 48 h), suggesting FMs accumulated mainly in pDCs (**Figure 3B** and representative gating strategy shown in **Figure S2**). Together, these results demonstrated the highly targeted accumulation of FMs in pDCs despite their relatively low abundance.

**Figure 3.**
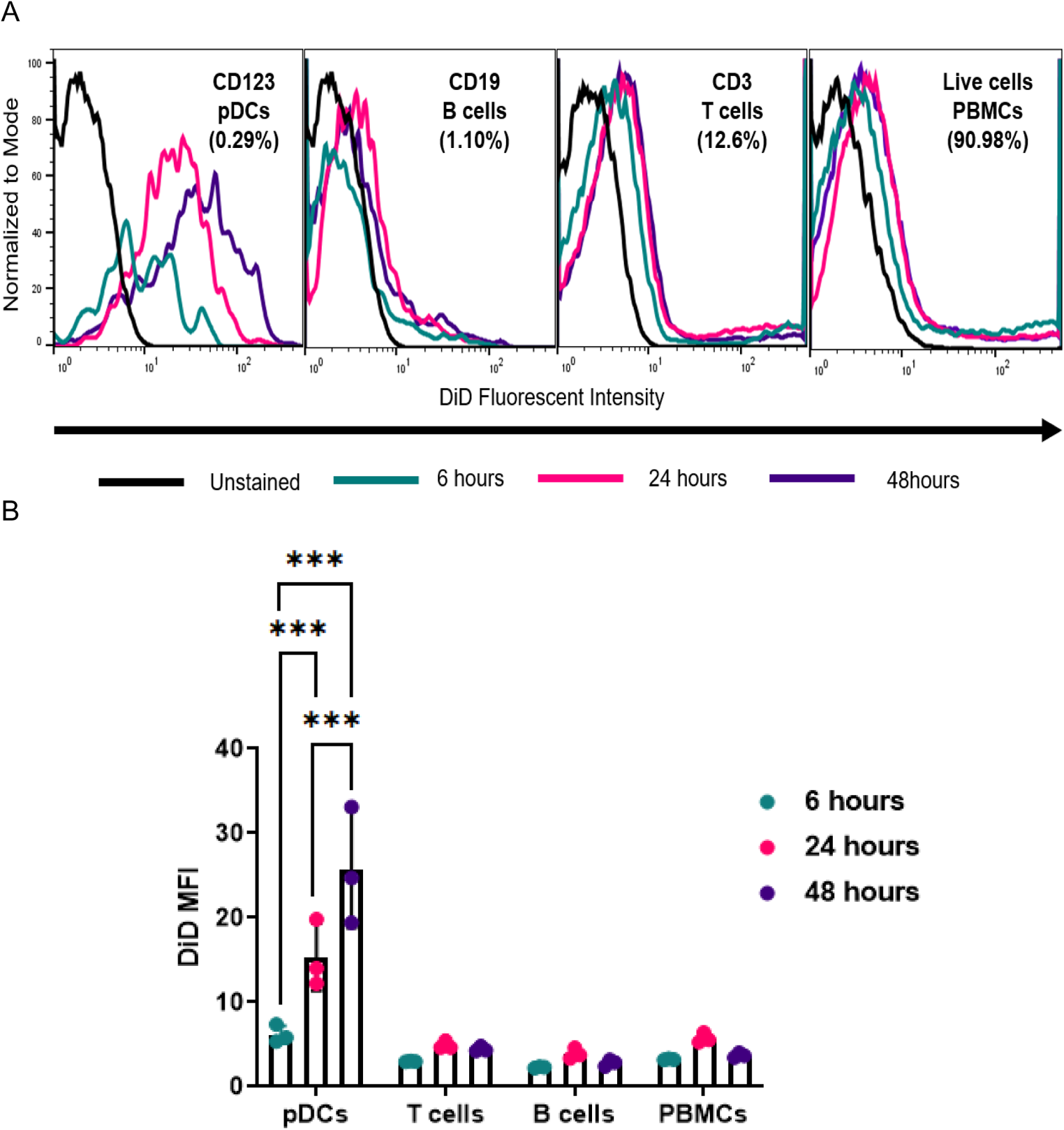
DiD-loaded FMs preferentially accumulated in pDCs *in vitro*. FMs were loaded with DiD fluorescent tracer dye. Fresh human PBMCs were cultured with 200 µg/mL DiD-loaded FMs for 6, 24 and 48 h in supplemented RPMI 1640 plus GlutaMAX medium with 10% FBS and 20 ng/mL IL-3 at 37°C and 5% CO_2_ (*n* = 3 independent donors). (A) Representative histograms and (B) median fluorescent intensity of DiD from (A). Bars represent mean (*n* = 3) with standard deviation. Statistical analysis: Tukey’s multiple comparison two-way ANOVA and *p* <0.05 was considered significant: ***p ≤ 0.001. Gating strategies are described in Figure S2.

The biodistribution of nanocarriers is strongly influenced by parameters such as size, morphology, dose, and administration route. Typically, accumulation in blood filtration organs (*e.g.*, kidney, liver, spleen) is undesirable for drug delivery because nanocarriers are sequestered by the mononuclear phagocyte system (38, 41). In SLE, these are critical sites of disease activity for targeting drug delivery. Biodistribution of DiD-loaded FMs was analyzed after intravenous injection in C57BL/6 mice. FMs accumulated in the kidneys and liver 1-h post-injection and dye signal was cleared by the body after 24 h with minimal accumulation in the eye (**Figure 4A**). The decreased accumulation of FMs in the eye suggests the potential to reduce antimalarial retinopathy by minimizing off-target drug accumulation, potentially eliminating a primary toxicity of soluble CQ.

**Figure 4.**
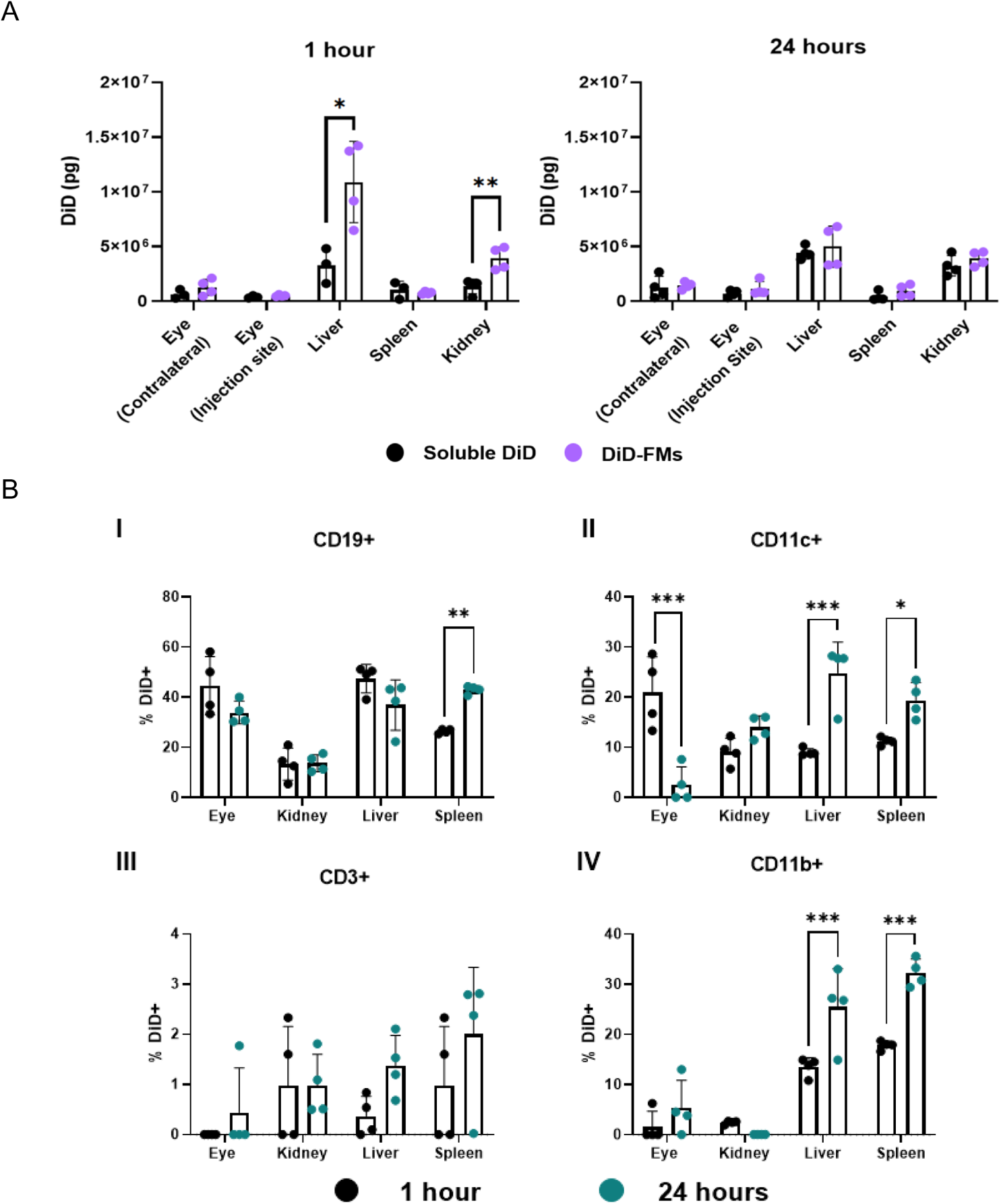
Organ- and cellular-level biodistribution of DiD-loaded FMs in C57BL/6 Mice. DiD-loaded FM were synthesized as described and administered by intravenous injection into the retro-orbital sinus. Mice were sacrificed at 1 h or 24 h post-injection and organs were mechanically dissociated and digested by collagenase, type 4. (A) The absorbance at 644 nm of 250,000 single cell suspensions from C57BL/6 mice (*n* = 3-4) at each organ was calculated against a standard curve of DiD in 1X DPBS. Graphs compare tissue distribution of DiD-loaded FMs or soluble FMs at both 50 µg/mL. Statistical analysis completed by unpaired t tests. Statistical significance: *p ≤ 0.05 and **p ≤ 0.01. (B) Single-cell suspensions were stained and analyzed by flow cytometry. Immune cells from both eyes were pooled, stained, and analyzed by flow cytometry. Graphs compare cellular uptake of DiD-loaded FMs at different time points. Each cell type was mutually exclusive to other cell markers. Particle uptake was analyzed in the following cells: (I) CD19+ B cells, (II) CD11c+ dendritic cells, (III) CD3+ T cells, and (IV) CD11b+ myeloid cells. Statistical analysis was completed by two-way ANOVA with Šídák’s multiple comparison test. Statistical significance: *p ≤ 0.05, **p ≤ 0.01, and ***p ≤ 0.001.

Within each organ, we analyzed the percentage of immune cells that were DiD+ to determine FM uptake (**Figure 4B**). We observed a significant increase in splenic CD19+ B cells associated with DiD-loaded FMs after 24 h. CD11c+ dendritic cells and CD11b+ myeloid cells also showed increased DiD signal between 1- and 24-h post-injection in the liver and spleen, consistent with their passive endocytic function. This may also be partially driven by the high aspect ratio and minimal curvature regions of FMs (normalized curvature, Ω ≥ 45°), which can induce faster internalization by phagocytosis compared to spherical nanocarriers (Ω ≤ 45°) (42). The increased cellular signal from 1 h to 24 h may be the result of cells in the mononuclear phagocyte system facilitating the clearance of FMs from circulation and at those organ sites. These results demonstrated favorable drug delivery properties at the tissue and cellular level to address key disadvantages of a soluble free drug, including avoiding the eye and targeting drugs into immunopathogenic cell types and tissue sites of disease activity.

### Chloroquine-Loaded Nanocarriers Decrease Interferon-Stimulated Genes in Human PBMCs

A major goal for targeted delivery and controlled release is to potentiate drug activity. We compared the efficacy of CQ-loaded nanocarriers to soluble CQ in two *in vitro* culture systems: human PBMCs stimulated with 1) TLR agonists or 2) sera from SLE patients with active disease. We generated a dose-response curve to determine which concentrations of CQ inhibited TLR7 and TLR9 activation in healthy human PBMCs. Previous studies show that 100 µM CQ maximally decreases pro-inflammatory cytokine secretion, including TNF-α, IL-6, and IL-1β (43, 44), and particularly, IFN-α in human PBMCs after *in vitro* TLR stimulation (45). We quantified the expression of *MX1*, an interferon-stimulated gene that is upregulated in SLE patients compared to healthy controls (46). Healthy human PBMCs were stimulated with either ssRNA40/LyoVec (TLR7/8 agonist) or CpG ODN 2216 Class-A (TLR9 agonist) and left untreated or pretreated with soluble drug or drug-loaded nanocarriers at 1.95, 3.91, or 7.81 μM CQ (**Figure 5**). CQ-loaded FMs resulted in >90% decrease in *MX1* gene expression in human PBMCs after TLR7/8 (**Figure 5A**) and TLR9 (**Figure 5B**) activation.

**Figure 5.**
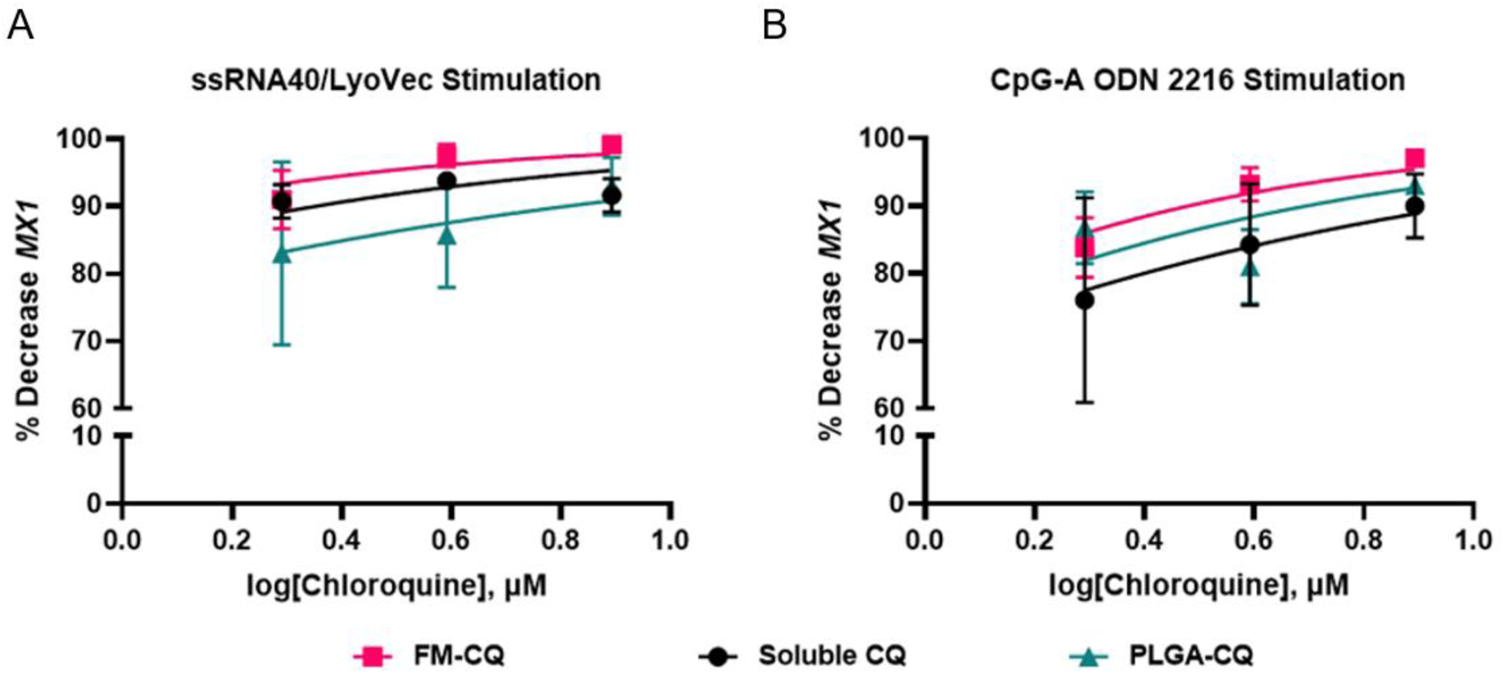
Dose-response curves of CQ concentration versus percent decrease in *MX1* gene expression. Human PBMCs (*n* = 4 independent donors) were pretreated with CQ or CQ-loaded FM or PLGA nanocarriers for 1 h and then stimulated with (A) ssRNA40/LyoVec (TLR7/8 agonist) for 4 h or (B) CpG-A ODN 2216 (TLR9 agonist) for 6 h. RNA was extracted to quantify *MX1* expression using TaqMan real-time RT-qPCR and normalized to β-actin. *MX1* expression was normalized to TLR agonist alone. Lines represented the non-linear fit of log(inhibitor) versus normalized response - variable slope model, where CQ was the inhibitor concentration. Data are means ± standard deviation.

Using the dose-response curve data, we subsequently tested CQ-loaded nanocarriers at 3.91 μM CQ because this dose yielded robust inhibition of *MX1* (93.20-97.57%) at approximately 25-fold lower dose than previous studies with soluble drug (43–45) and less variability in response after both TLR7/8 and TL9 activation. Soluble CQ and CQ-loaded nanocarriers had comparable efficacy suppressing TLR7/8-mediated *MX1* gene expression in PBMCs (**Figure 6A**). In contrast, CQ-loaded FMs were significantly more suppressive of TLR9-mediated *MX1* expression in PBMCs, approximately 2.6-fold more inhibitory than soluble drug (**Figure 6B**). Nanocarriers alone, only soluble CQ, or blank nanocarriers cultured with soluble CQ did not stimulate *MX1* gene expression in healthy human PBMCs (**Figure S3**).

**Figure 6.**
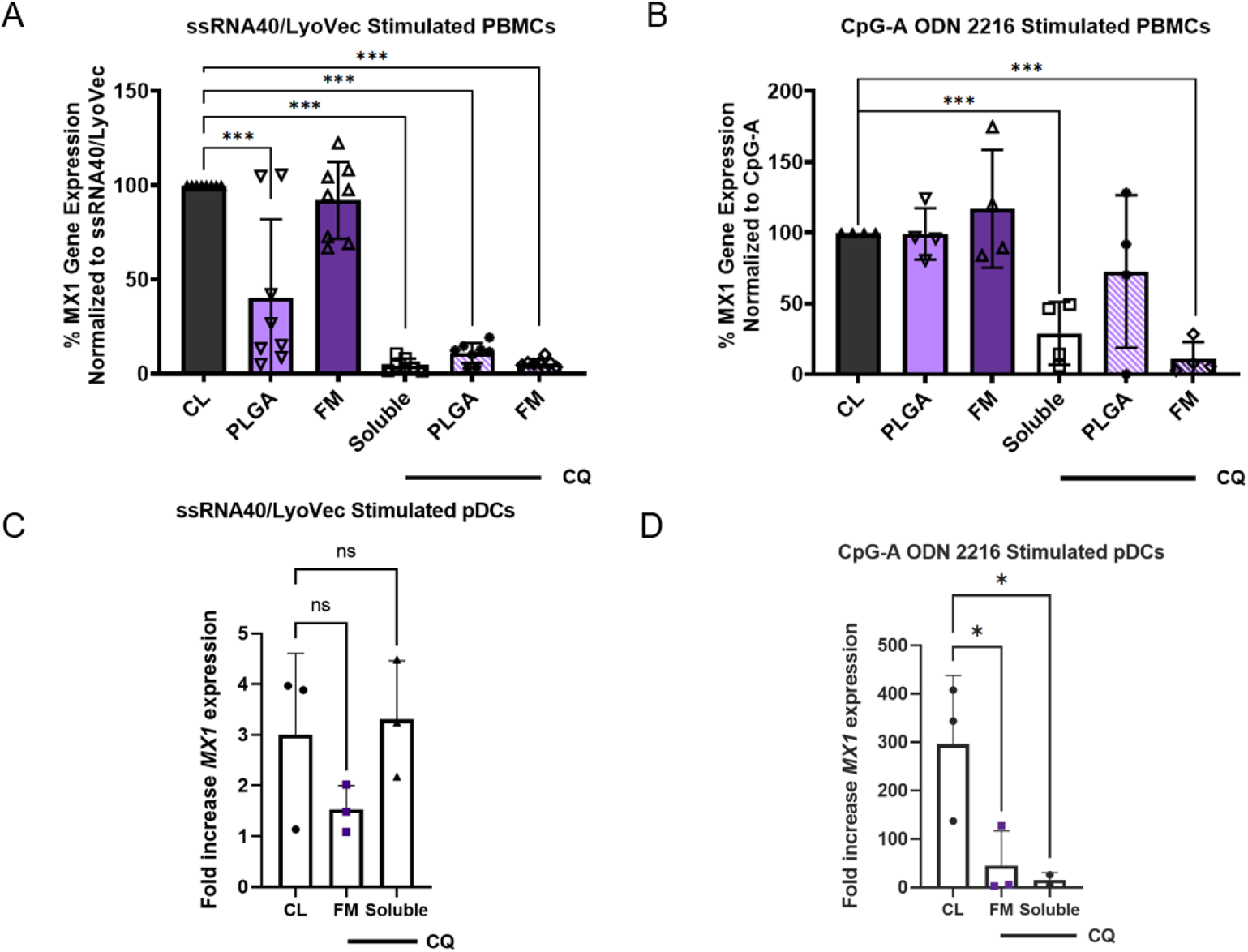
Soluble and nanocarrier-delivered CQ decreased *MX1* expression in TLR-stimulated human PBMCs and isolated pDCs. Healthy human PBMCs were isolated, treated with 3.91 μM soluble or encapsulated CQ or equal mass blank nanocarriers for 1 h, and then activated for 4 h with (A) 5 μg/mL ssRNA40/LyoVec or for 6 h with (B) 5 μM CpG A 2216. Total RNA was isolated and *MX1* expression quantified using TaqMan RT-qPCR normalized to β-actin expression. All samples were normalized to TLR agonists alone. Data are means ± standard deviation for *n* = 4-8 independent donors. Statistical significance was evaluated by one-way ANOVA with Dunnett’s multiple comparisons test, ***p-adjusted ≤ 0.001 and ****p ≤ 0.0001. PBMCs were processed by magnetic bead negative selection to purify pDCs. Isolated human CD123+ pDCs at 100,000 cells per condition (*n* = 3 independent donors) were pre-treated with soluble or CQ loaded FMs at 3.91 μM for 1 h and then stimulated with (C) 5 µg/mL ssRNA40/LyoVec or (D) 5 µM CpG-A for 4 or 6 h, respectively. Total RNA was isolated to analyze *MX1* expression using TaqMan assays with β-actin control. Statistical analysis: one-way ANOVA with Dunnett’s multiple comparisons test where \**p* ≤ 0.05. Abbreviation: CL, control.

pDCs are well-known as the primary producers of type I IFN among immune cells in PBMCs, but they are notoriously low in abundance. We tested the contribution of pDCs to TLR-driven type I IFN by isolating purified human CD123+ pDCs, stimulating them with TLR agonists, and treating them with either soluble CQ or CQ-loaded nanocarriers (**Figure 6C** and **6D**). Purified pDCs showed no significant *MX1* upregulation after activation by TLR7/8 agonist ssRNA40/LyoVec (**Figure 6C**). This suggested the responding PBMC cell type to TLR7/8 stimulation was not pDCs in our experiments. Unlike TLR7/8 stimulation, TLR9 stimulation of pDCs resulted in robust *MX1* upregulation, which could be strongly inhibited by both soluble CQ and CQ-loaded FMs. TLR9 stimulation with CpG-A ODN 2216 robustly stimulated type I IFN in purified pDCs, upregulating *MX1* expression ~300X over unstimulated controls after 6 h (**Figure 6D**). Soluble CQ and CQ-loaded FMs significantly decreased *MX1* (95% and 85% inhibition, respectively). Overall, these results demonstrated that soluble CQ and CQ-loaded nanocarriers can efficiently suppress TLR7/8 and TLR9-stimulated type I IFN responses in PBMCs, and that pDCs were a primary target in TLR9 stimulation while a different PBMC cell type drove TLR7/8 responses.

### Chloroquine Loaded Nanocarriers Reduce Type I IFN Induced by SLE Sera

Purified TLR agonists are strong stimulators of PBMCs but differ substantially from physiologic agonists. Circulating immune complexes in SLE are unique structures composed of autoantibodies and endogenous antigens, such as anti-dsDNA antibodies and self-DNA, and these are hypothesized to be a major driver of endosomal TLR activation and type I IFN pathogenesis in SLE. Anti-dsDNA autoantibodies are found in approximately 80% of patients with lupus nephritis (47), and are associated with TLR9 activation (48), and high IFN-α activity (49). SLE serum has been previously shown to stimulate pDCs and produce IFN-α (50). CQ also has been shown to decrease IFN-α production after pDC activation by SLE serum (19). We used SLE patient sera as a more physiologic stimulator of PBMCs and proof-of-principle for clinical utility. Healthy PBMCs were isolated and co-cultured with 30% v/v SLE sera for 24 h *in vitro*. We used a ~75% lower dose of soluble CQ than reported in literature to inhibit SLE serum (19). Soluble CQ did not significantly decrease type I IFN response stimulated by SLE sera (**Figure 7A**). In contrast, pretreatment with equivalent dose of CQ-loaded FMs significantly decreased *MX1* expression induced by SLE sera by approximately 75% compared to no inhibition by either soluble drug or CQ-loaded spherical PLGA nanocarriers (**Figure 7A**). We also analyzed secretion of IFN-α with a multi-subtype ELISA that quantifies all 12 known IFN-α family proteins in humans (51). As expected, based on *MX1* expression, IFN-α production was significantly decreased by CQ-loaded nanocarriers, while soluble CQ had no effect (**Figure 7B**). Both FMs and spherical PLGA nanocarriers significantly inhibited IFN-α secretion, suggesting *MX1* is induced by more than IFN-α in our experimental system. These data showed that CQ-loaded nanocarriers significantly decreased IFN-α production compared to soluble CQ in response to immune complexes from SLE patient sera. This suggests that CQ-loaded FMs may have utility as a novel drug formulation for treating SLE.

**Figure 7.**
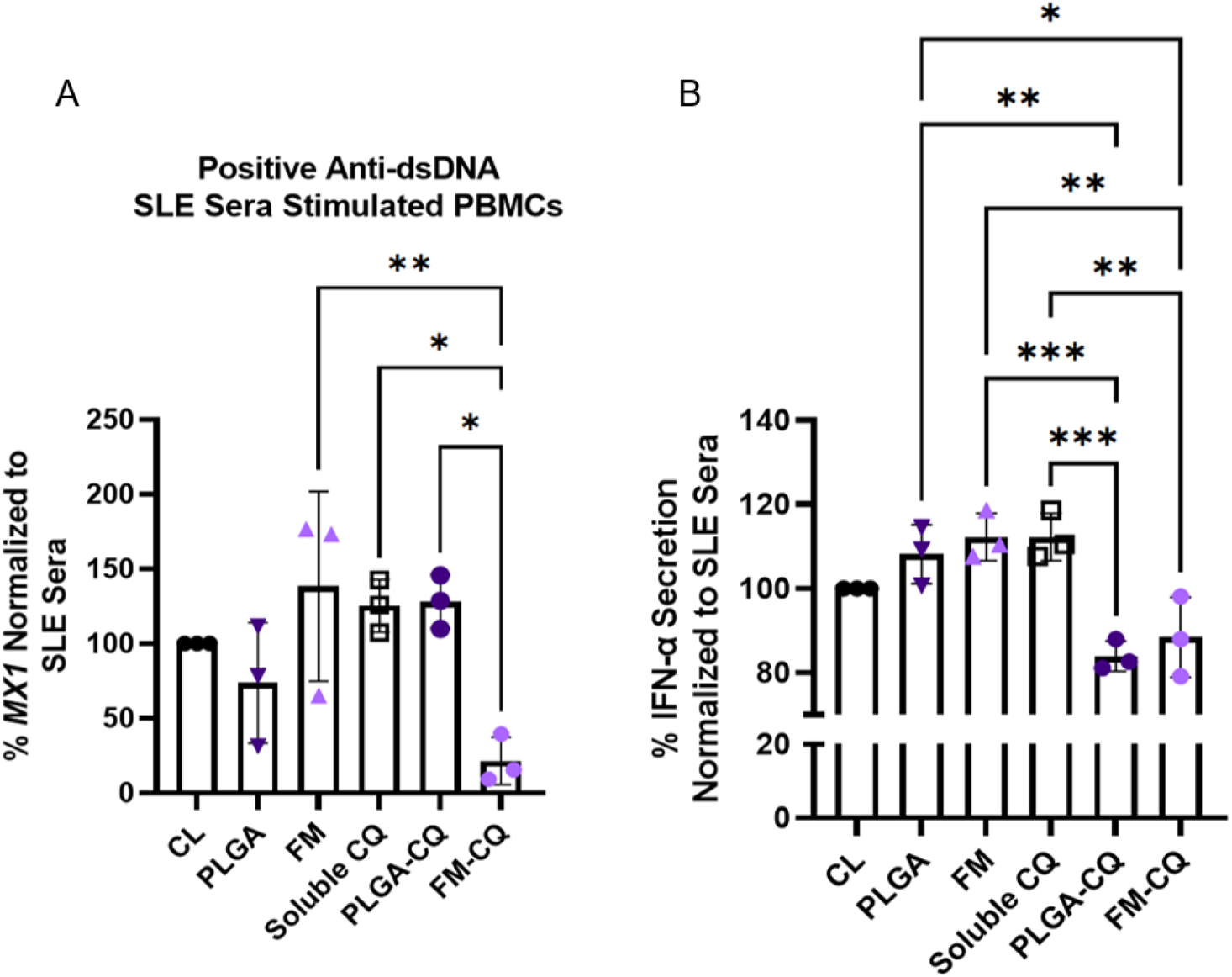
CQ-loaded nanocarriers suppressed SLE sera-induced type I IFN responses from human PBMCs. Healthy human PBMCs were treated with soluble or CQ-loaded nanocarriers at 3.91 μM or equal mass blank nanocarriers for 1 h, then stimulated with 30% v/v SLE patient sera for 24 h. (A) Total RNA was isolated and *MX1* expression quantified using TaqMan RT-qPCR normalized to β-actin expression. (B) Supernatants from cell culture were collected for multi-subtype IFN-α ELISA. All samples were normalized to SLE sera alone. Data are means ± standard deviation for *n* = 3 healthy independent PBMC donors. Statistical significance was evaluated by one-way ANOVA with Tukey’s multiple comparisons test, such that, *p ≤ 0.05, **p ≤ 0.01, and ***p ≤ 0.001. Abbreviation: CL, control.

## Discussion

The past decade has seen the first new drug approvals for SLE by the U.S. FDA in over 50 years. In 2011, the monoclonal antibody belimumab blocking B lymphocyte stimulator was approved for SLE and approved for lupus nephritis in 2020. In 2021, voclosporin, a calcineurin inhibitor, in combination with background immunosuppressive therapy, was approved for adult patients with active lupus nephritis. Voclosporin was the second FDA-approved therapy for lupus nephritis and the first oral treatment specifically for that manifestation (52). Despite FDA approvals of belimumab and voclosporin, these therapies were studied in combination with a background of immunosuppressive agents and corticosteroids to achieve efficacy, and drug delivery and toxicity remain persistent limitations for these and other SLE treatments.

Nanocarriers can enhance drug delivery while mitigating adverse side effects by reducing accumulation in off-target cells and tissues. We demonstrated that targeting pDCs using antimalarial-loaded FMs could enhance suppression of TLR activation and subsequent type I IFN responses (**Figure 1**). To our knowledge, this is the first use of passive, morphology-based nanocarrier targeting of pDCs to enhance TLR inhibition, and extends prior work demonstrating accumulation in pDCs (33). TLR inhibition has great potential as a therapeutic strategy since gene polymorphisms of TLRs lead to disease susceptibility (53), TLR ligands contained in NETs or resulting from apoptotic cell death exacerbate disease (54, 55), and TLR inhibitors (*i.e*., antimalarial drugs) alleviate disease (56). Our approach targets SLE in disease-relevant sites and cells to inhibit key inflammatory pathways.

Our results showed that FMs accumulate in pDCs (**Figure 3**), professional IFN-α producing cells that represent <1% of cells in the blood but produce 1,000X more IFN-α than any other immune cell (57). Concentrating drug delivery to pDCs may potentiate TLR antagonists, such as CQ, by increasing drug accumulation in the endosomal space to block TLR signaling more efficiently and downstream type I IFN responses. Additionally, passive targeting, which leverages physicochemical nanocarrier properties (**Figure 2**) and disease biology, presents an opportunity for FMs to accumulate at tissue sites of SLE-driven inflammatory damage like the kidneys (**Figure 4**). Targeting SLE-relevant organs (*e.g*., kidneys and liver) can concentrate drug delivery to tissues where both immune complexes and pDCs are found during active disease, while limiting off-target effects. pDCs are important in lupus nephritis because they express high levels of IL-18R which allows the relocation of dendritic cells within the glomeruli by IL-18 stimulation. These dendritic cells then activate resident T cells, resulting in promotion of renal damage (58, 59). FMs also escape accumulation in the eye (**Figure 4**) due to the size and morphology of the nanocarriers and their impermeability of the blood-retinal barrier (60, 61). This may reduce the risk of antimalarial-mediated retinopathy with chronic use of CQ (29) and long-term toxicity associated with the current CQ formulation. We hypothesize that the filamentous morphology (**Figure 2**) and phagocytosis of FMs facilitates internalization by pDCs (34, 42, 62), and thus, its unique morphology advances nanocarrier drug delivery. Future studies should evaluate the molecular mechanisms and kinetics of FM uptake into endosomes and degradation by pH-mediated oxidation.

We used synthetic oligonucleotide agonists ssRNA40 and CpG-A ODN 2216 to trigger TLR7/8 and TLR9, respectively, because human pDCs selectively express endosomal TLR7 and TLR9 to sense pathogenic and endogenous nucleic acids (63). Although ssRNA40 is known to produce high levels of IFN-α in pDCs (64) and induce *MX1* gene expression in PBMCs (65), CQ-loaded FMs were not significantly different from soluble drug or spherical PLGA control (**Figure 6A** and **6C**). Since isolated pDCs did not show an *MX1* response to ssRNA40, we hypothesize that monocytes and other myeloid cells may be additional mediators of IFN-α production after TLR7/8 activation by ssRNA40 (66). However, both soluble CQ and CQ-loaded FMs did suppress TLR9-mediated *MX1* in both PBMCs and isolated pDCs at the same CQ concentration (**Figure 6B** and **6D**). This represents a major advantage in that the total body exposure and off-target tissue accumulation of CQ is diminished with the delivery of antimalarial drug in nanocarriers.

Many emerging drug delivery strategies have been tested preclinically against SLE. Previous studies have used mycophenolic acid (67, 68), cyclosporine A (69), azathioprine (70), and the prodrug of dexamethasone (71) in drug delivery platforms to target SLE relevant pathways, ameliorate glomerulonephritis in lupus-prone mice, and decrease proinflammatory cytokines. Our work builds upon existing studies by using a novel drug delivery approach to target pDCs without needing a targeting moiety for delivery of an FDA-approved drug, CQ, for SLE treatment. We chose antimalarial drugs because they are the mainstay first line, long-term SLE treatment regardless of renal involvement or disease severity (25). By loading CQ in FMs, we decreased the *in vitro* concentration of CQ required to inhibit *MX1*, which is upregulated in SLE patients (72) (**Figure 5**). We showed that CQ-loaded FMs decreased *MX1* gene expression equivalent to soluble CQ in human PBMCs stimulated with purified TLR agonists (**Figure 6A** and **6B**). In human PBMCs stimulated with anti-dsDNA positive SLE sera, soluble CQ did not inhibit *MX1* gene expression (**Figure 7A**) or IFN-α secretion (**Figure 7B**). We propose that SLE sera is less sensitive to CQ inhibition than synthetic oligonucleotide TLR agonists, as shown in previous studies (19), and concentration of CQ in the endosomal space may be more important for treatment in this scenario. Compared to previous studies, we used 75% less CQ loaded in FMs and showed a significant decrease in *MX1* gene expression (**Figure 7A**) and IFN-α production (**Figure 7B**), demonstrating the dose-sparing and dose-enhancement of CQ in FMs versus soluble CQ.

Future studies evaluating the dosing strategies of CQ loaded nanocarriers in lupus-prone mouse models can better define the frequency of treatment as well as determine bioavailability of different routes of administration. A preclinical model with a strong type I IFN signature and clinical manifestations similar to SLE patients, such as the pristane-induced model (73, 74), will be important in demonstrating the efficacy of this treatment option *in vivo*. Our approach has therapeutic implications because CQ-loaded FMs may provide a more targeted inhibition of immune-complex-mediated inflammation in SLE, potentially sparing steroid or immunosuppressive treatment. This would result in both lower steroid toxicity as well as lower risk of infection, a serious threat to SLE patient health (15, 16).

To enhance the therapeutic potential of CQ drug delivery, future work should include investigating other nanocarrier morphologies that can target other IFN-producing cell types such as myeloid cells. The PEG-*b*-PPS platform is ideal for these studies because by changing the hydrophilic weight fractions of the polymer, they can be self-assembled easily to form diverse morphologies such as spherical (*i.e.*, micelles and polymersomes) (32) and cubic (*i.e*., bicontinuous cubic nanospheres) (75) nanostructures with distinct cellular biodistribution profiles (76). Overall, this study illustrates the therapeutic potential of drug delivery of CQ for targeting SLE-relevant pathways, immune cells, and organ sites.

## Methods

### Materials

Chloroquine, 95% purity was purchased from Ark Pharm (Arlington Heights, IL, USA). Acid-terminated, 50:50 lactide/glycolide molar ratio, molecular weight (MW) 7,000-17,000 poly(D, L-lactide-co-glycolide) (PLGA); MW 25,000, 88% hydrolyzed polyvinyl alcohol (PVA); poly(ethylene glycol) methyl ether MW 2000; and organic solvents were purchased from Sigma Aldrich (St. Louis, MO, USA). Propylene sulfide was acquired from TCI Chemicals. Micro Float-A-Lyzer Dialysis Device, biotech grade cellulose ester, 8-10 kDa molecular weight cut off (MWCO) was purchased from Repligen (Waltham, MA, USA). TLR agonists (ssRNA40/LyoVec and CpG-A ODN 2216) were purchased from Invivogen (San Diego, CA, USA).

### Antibodies and Dyes

The Zombie Aqua™ Fixable Viability dye and all primary antibodies used for flow cytometry analysis were obtained from BioLegend (San Diego, CA, USA) (**Table 1**).

**Table 1.**
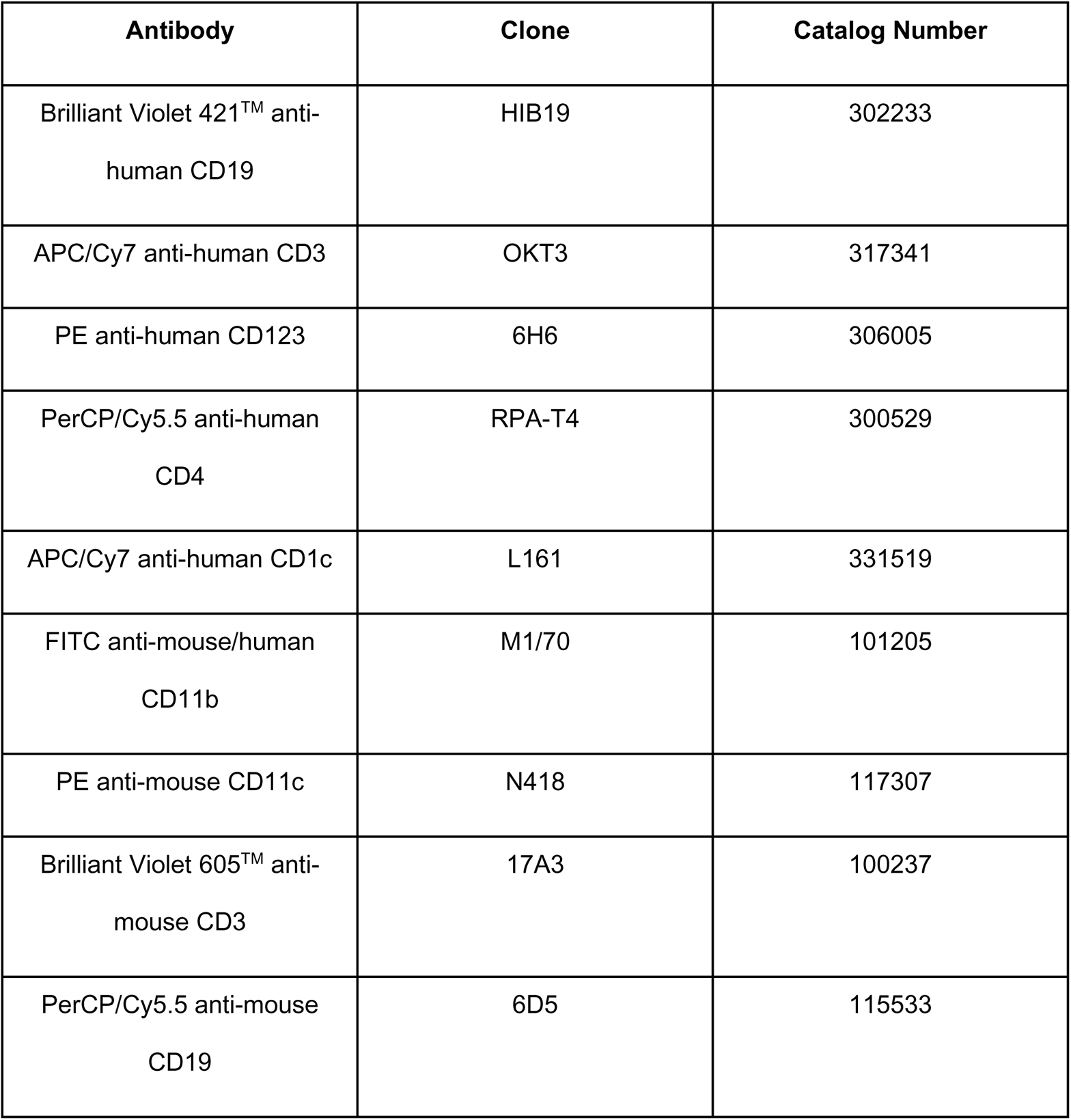
Flow Cytometry Antibodies

### Nanocarrier Synthesis

Block copolymer PEG-*b*-PPS was synthesized as previously described (77). FMs were loaded with the hydrophobic antimalarial drug, CQ, via thin-film hydration. Briefly, 5-10 mg of PEG_45_-*b*-PPS_44_ polymer was co-dissolved with equal mass CQ in 200 µL chloroform in a 5 mL sterile, clear, LPS-free glass vial. The solvent was evaporated at room temperature (RT) for 3-5 h, resulting in a thin film. The thin film was rehydrated at a total polymer concentration of 5-15 mg/mL with 1X phosphate-buffered saline (PBS) at pH 7.4 under gentle agitation overnight. CQ loaded nanocarriers were purified from free CQ by 10K MWCO polyethersulfone membrane spin columns (VWR; Radnor, PA, USA) at 10,000 xg for 1-2 minutes and equilibrated with PBS solution.

Spherical PLGA nanocarriers were fabricated by oil-in-water emulsion/solvent evaporation technique (78). Briefly, 5-10 mg PLGA and equal mass CQ was dissolved in 500 µL chloroform. The organic phase was emulsified in 2 mL of 0.5% PVA in 1X PBS solution using a probe sonicator at 50% amplification for 2 minutes on ice. The emulsion was added dropwise to 10 mL of 5% PVA in 1X PBS solution at RT. The continuous phase was homogenized at 6,800 x rpm with the T 25 digital ULTRA-TURRAX® (IKA; Wilmington, NC, USA) and left to stir overnight at 600 x rpm at RT until the organic solvent was evaporated and nanocarriers were hardened. CQ-loaded spherical PLGA nanocarriers were purified from free drug by centrifugation for 5 minutes at 14,000 x g and washed three times with diH_2_O.

### Nanocarrier Characterization

Small-angle X-ray scattering (SAXS) was performed at the University of Maryland, College Park X-Ray Crystallographic Center using the Xenocs Xeuss SAXS/WAXS/GISAXS small-angle system with 8 keV (wavelength = 1.5 A) collimated X-rays. Samples were measured at 2.5 m from the CCD detector and analyzed within the 0.004-0.2 A^−1^ q-range calibrated by diffraction patterns of silver behenate. SAXS analysis was performed using IgorPro (WaveMetrics, Inc.; Portland, OR, USA) for 2D to 1D reduction and normalization of acquired sample scattering from buffer scattering. Model fitting was completed using SASVIEW 4.X based on the flexible cylinder model with the following parameters: 2 µm cylinder length, 150 nm persistence length, and 8 nm PPS core radius (79). The FEI Morgagni 268 100 kV Transmission electron microscope (TEM) equipped with a Gatan Orius CCD camera was used to compare morphology and appearance of blank and CQ-loaded nanocarriers. For TEM processing, 12 mg/mL blank or loaded nanocarriers were added to FCF200-Cu-TB coated grids and stained with 2% uranyl acetate. The grids were immediately processed after negative staining. Size, size distribution, and zeta potential were measured at 1 mg/mL of nanocarriers in 1X PBS solution (pH 7.4) using the Malvern Zetasizer Nano ZS. Loading capacity, total loaded drug per mass of nanocarrier, and encapsulation efficiency, percentage of initial drug mass successfully entrapped in the nanocarriers, were measured by dissolving a known mass of loaded nanocarriers in dimethyl sulfoxide (DMSO) followed by UV/Vis spectrophotometry analysis using the Mettler Toledo UV5 Nano. Drug loading per mass of nanocarriers was determined by developing a standard curve of CQ dissolved in DMSO using the characteristic wavelength corresponding to maximum absorption of CQ at approximately 343 nm. Drug release kinetics were determined by placing drug-loaded nanocarriers into microdialysis devices in 1X PBS with 1% bovine serum albumin (BSA) at 37 °C (physiological temperature) and 5% CO_2_. Samples were analyzed by UV/Vis spectrophotometry, as above, after 0, 2, 4, 6, and 24 h.

### Nanocarrier Cellular Uptake and In Vivo Biodistribution

FMs were loaded with the lipophilic fluorescent tracer, Vybrant® DiD cell-labeling solution (Thermo Fisher; Waltham, MA, USA), via thin-film hydration method. In a 5 mL sterile, clear, LPS-free glass vial, 2.5 µL Vybrant® DiD cell-labeling solution and 5-10 mg of PEG-*b*-PPS was co-dissolved in 150 µL dichloromethane. The solvent was evaporated at RT for 3-5 h and then resuspended at a total polymer concentration of 5-15 mg/mL with 1X PBS at pH 7.4 under gentle agitation overnight. Dye-loaded nanocarriers were purified from free dye by dialysis or 10K MWCO polyethersulfone membrane spin columns (VWR; Radnor, PA, USA) at 10,000 xg for 1-2 minutes and equilibrated with PBS solution. Lipophilic dye loaded nanocarriers at 200 µg/mL were incubated at 37°C, 5% CO_2_ in 1 million fresh human peripheral blood mononuclear cells (PBMCs) from healthy donors (New York Blood Center; New York, NY) for up to 48 h in RPMI 1640 medium with GlutaMAX plus 10% fetal bovine serum (FBS), 1% sodium pyruvate, 1% nonessential amino acids (NEAA), 1% penicillin-streptomycin solution (pen-strep), and 20 ng/mL recombinant human IL-3 (BioLegend; San Diego, CA, USA) to determine cellular distribution. PBMCs were stained with a viability dye and fluorescent antibodies to distinguish cellular subsets including CD19+ B cells (clone HIB19; Brilliant Violet 421™), CD3+ T cells (clone OKT3; APC/Cy7), and CD123+ pDCs (clone 6H6; PE).

Female C57BL/6 mice, 6-8 weeks old, (Jackson Laboratory; Bar Harbor, ME USA) were injected intravenously by retro-orbital sinus with 150 μL free DiD (50 μg/mL in 1X PBS solution) or DiD loaded FMs (7.5 μg of loaded DiD in each sample in 150 μL PBS). At 1- and 24-h post-injection, the eyes, kidneys, liver, and spleen were collected, mechanically dissociated, and digested with 0.1% collagenase, type 4 in Hanks’ Balanced Salt Solution (Worthington; Lakewood, NJ, USA) followed by red blood cell lysis with ammonium-chloride-potassium lysing buffer (Lonza; Basel, Switzerland). Cells were filtered through a 70 μm nylon, DNase/RNase free, non-pyrogenic cell strainer (VWR; Radnor, PA, USA) and washed with 1X PBS. Single cell suspensions of each tissue at 250,000 cells per 200 μL in 1X DPBS were read on a BioTek Cytation 5 plate reader at the absorbance maximum of DiD (*i.e*., 644 nm). The total mass of DiD per tissue was quantified by interpolation of sample measurements onto a standard curve of known concentrations of DiD in 1X DPBS using the characteristic maximum absorbance. Cell suspensions were stained for viability and phenotypic, anti-mouse cell surface markers, CD11b (clone: M1/70, FITC), CD11c (clone: N418, PE), CD3 (clone: 17A3, Brilliant Violet 605™), and CD19 (clone: 6D5, PerCP/Cy5.5). Stained cells were then run on a flow cytometer and analyzed for uptake of DiD loaded nanocarriers.

### In Vitro Activity of Nanocarriers in Human Cells

Fresh, healthy human PBMCs were isolated from buffy coats obtained from the New York Blood Center via density gradient medium (Ficoll-Paque). Flow cytometry was used to establish viability and cell distribution of each human PBMC donor using the following fluorophore-conjugated anti-human antibodies: CD4+ T cells (clone RPA-T4; PerCP-Cy5.5), CD19+ B cells (clone HIB19; Brilliant Violet 421™), CD123+ pDCs (clone 6H6; PE), and CD1c+ mDCs (clone L161; APC/Cy7). One million PBMCs in 250 μL total volume of RPMI 1640 with GlutaMAX medium plus 10% FBS, 1% sodium pyruvate, 1% NEAA, and 1% pen-strep were treated with soluble CQ, empty nanocarriers, or CQ-loaded nanocarriers for 1 h and then stimulated with purified TLR agonists: 5 μg/mL ssRNA40/LyoVec (TLR7/8 agonist) (64), or 5 μM CpG-A ODN2216 (TLR9 agonist) (80), or 30% v/v sera from an SLE patient with moderately active disease and positive anti-dsDNA titers for 4, 6, and 24 h, respectively. Recognition of ssRNA40 is species-dependent where human cells have a bias towards TLR8 and mouse cells bias towards TLR7 (66). Chloroquine was tested at 1.95, 3.91, or 7.81 μM. To prevent degradation and enhance cellular uptake of the TLR7/8 agonist, RNA was co-delivered using LyoVec as a transfection agent. Following incubation at 37 °C, 5% CO_2_, cells were spun down at 500 xg for 1 minute at RT. Total RNA was isolated from PBMCs using the Quick-RNA Miniprep Plus Kit (ZymoResearch; Irvine, CA, USA), amplified, and analyzed by real-time reverse transcription-quantitative PCR (RT-qPCR). Cell-free supernatants were collected and stored at −80°C until testing for IFN-α production using the Human IFN Alpha All Subtype ELISA Kit (PBL Science; Piscataway, NJ, USA). Nanocarrier efficacy was determined by analyzing the downstream IFN-stimulated gene *MX1* by RT-qPCR. Expression levels were normalized to β-actin control. Purified pDCs were isolated via the EasySep™ Human Plasmacytoid DC Enrichment Kit (STEMCELL Technologies; Vancouver, Canada). Soluble or CQ-loaded nanocarriers at 3.91 μM total drug were cultured with 100,000 pDCs in RPMI 1640 with GlutaMAX plus 10% FBS, 1% sodium pyruvate, 1% NEAA, 1% pen-strep, and 20 ng/mL recombinant human IL-3. Cells and cell-free supernatants were isolated from stimulated pDC cultures and tested by real-time RT-qPCR and human IFN-α all subtype ELISA, respectively.

### Flow Cytometry

Either Human TruStain FcX™ or TruStain FcX™ PLUS (BioLegend; San Diego, CA, USA) was used to block nonspecific binding of human or mouse Fc receptors, respectively, prior to immunostaining. For human cells, 5 μL of Human TruStain FcX™ was added per million cells in 100 μL staining volume of PBS plus 1% BSA. Cells were incubated for 5 minutes at RT. For mouse cells, 0.25 µg of TruStain FcX™ PLUS was added per million cells in a volume of 100 µL PBS plus 1% BSA for 5 minutes at RT. After blocking, cells were stained at 1:100 dilution with conjugated fluorescent antibodies (**Table 1**) in PBS plus 1% BSA for 15-20 minutes in the dark and on ice. Cells were washed with PBS. To discriminate between live and dead cells, Zombie Aqua™ Fixable Viability dye was diluted 1:1000 in PBS. Diluted Zombie Aqua™ Fixable Viability dye was added to cells for 15-30 minutes at RT and in the dark. Cells were washed, resuspended in PBS plus 1% BSA, and immediately analyzed on a flow cytometer. A BD LSR II with 405, 488, 561, and 640 nm excitation laser lines or Beckman Coulter CyAn ADP consisting of 405, 488, and 635 nm excitation laser lines was used for flow cytometry analysis of fluorescently labeled cells. Data were analyzed using FlowJo LLC software v10.7 (BD). The gating strategies are available in the supplementary figures.

### Real-time RT-qPCR

All real-time RT-qPCR reagents were purchased from ThermoFisher. The TaqMan probes included: Human MX1, FAM-MGB (assay id: Hs00895608_m1) and Human ACTB VIC-MGB PL (assay id: Hs01060665_g1). TaqPath 1-Step Multiplex Master Mix (No ROX) was used for all one-step multiplex real-time RT-qPCRs. Reactions were run on the CFX96 Touch Real-Time PCR Detection System using the following thermal cycle conditions: UNG incubation (1 cycle, 25°C, 2 minutes), reverse transcription (1 cycle, 53°C, 10 minutes), polymerase activation (1 cycle, 95°C, 2 minutes), and amplification (45 cycles at 95°C for 15 seconds and 60°C for 1 minute). The 2^−ΔΔCT^ (Livak) Method was used for relative gene expression analysis.

### Statistics

All statistical analyses were performed using GraphPad Prism 9 software (GraphPad Software, San Diego, CA). A minimum of three independent replicates were conducted for human PBMCs and nanocarrier characterization experiments. For *in vivo* biodistribution experiments, 3-4 mice were used. Paired or unpaired t-tests and ANOVA were used to test for statistical significance. P-values were adjusted for multiple comparisons by Tukey’s, Šídák’s, or Dunnett’s test, and adjusted p-values <0.05 were considered significant.

### Study approval

The animal study was reviewed and approved by UMBC Institutional Animal Care and Use Committee (OLAW Animal Welfare Assurance D16-00462). This study involved human subjects. Approval for this study was obtained from the University of Maryland School of Medicine Institutional Review Board (IRB) as well as the Baltimore VA Research Office of Human Research Protection. There is no identifiable medical information in this manuscript. All patient identities have been removed. Per our IRB-approved protocol, all participants signed informed consent. All identifiable information has been removed from the reported data.

## Author contributions

MEA and GLS conceptualized the study, designed experiments, analyzed the data, and wrote the manuscript. AG and VR contributed SLE patient sera and expertise in autoimmunity, rheumatology, and systemic lupus erythematosus. EAS contributed expertise in nanocarriers and drug delivery. MEA performed all experiments. EAS, NBK, SL, CJL, and SB synthesized the PEG-*b*-PPS polymer. All authors reviewed and edited the manuscript.

## Acknowledgments

We would like to thank Tagide deCarvalho for TEM assistance at the University of Maryland, Baltimore County Keith R. Porter Imaging Facility and Wonseok Hwang for SAXS assistance at the University of Maryland, College Park X-ray Crystallographic Center. Flow cytometry was performed at the University of Maryland Greenebaum Comprehensive Cancer Center Flow Cytometry Shared Service or the University of Maryland, Baltimore County Keith R. Porter Imaging Facility. We would like to thank Christine Daniel and Erin Lavik for the use of the ZetaSizer. This work was partially supported by the Lupus Foundation of America Gina M. Finzi Memorial Student Summer Fellowship (MEA), a grant from the State of Maryland, TEDCO Maryland Innovation Initiative (MII) (project #0719-009) (GLS), and the University of Maryland, Baltimore County Technology Catalyst Fund (GLS). AG wishes to acknowledge his completed VA CDA-2 award support—VA grant IK2 CX000649-01A1. MEA was supported by an NIH-NIGMS Initiative for Maximizing Student Development Grant (grant no. R25-GM55036) and the National Science Foundation LSAMP BD Program (award no. 1500511). This manuscript has been released as a preprint at bioRxiv.

## References

1. Singh RR, and Yen EY. SLE mortality remains disproportionately high, despite improvements over the last decade. Lupus. 2018;27(10):1577–81.

2. McDonald G, et al. Female Bias in Systemic Lupus Erythematosus is Associated with the Differential Expression of X-Linked Toll-Like Receptor 8. Front Immunol. 2015;6:457.

3. Murphy G, and Isenberg D. Effect of gender on clinical presentation in systemic lupus erythematosus. Rheumatology (Oxford). 2013;52(12):2108–15.

4. Mohan C, and Putterman C. Genetics and pathogenesis of systemic lupus erythematosus and lupus nephritis. Nat Rev Nephrol. 2015;11(6):329–41.

5. Fava A, and Petri M. Systemic lupus erythematosus: Diagnosis and clinical management. J Autoimmun. 2019;96:1–13.

6. Al Sawah S, et al. SAT0423 Understanding Delay in Diagnosis, Access to Care and Satisfaction with Care in Lupus: Findings from a Cross-Sectional Online Survey in the United States. Ann Rheum Dis. 2015;74(Suppl 2):812-.

7. Crow MK. Advances in understanding the role of type I interferons in systemic lupus erythematosus. Curr Opin Rheumatol. 2014;26(5):467–74.

8. Chasset F, and Arnaud L. Targeting interferons and their pathways in systemic lupus erythematosus. Autoimmun Rev. 2018;17(1):44–52.

9. Rönnblom L, and Leonard D. Interferon pathway in SLE: one key to unlocking the mystery of the disease. Lupus Sci Med. 2019;6(1):e000270.

10. Yan B, et al. Dysfunctional CD4+,CD25+ regulatory T cells in untreated active systemic lupus erythematosus secondary to interferon-alpha-producing antigen-presenting cells. Arthritis Rheum. 2008;58(3):801–12.

11. Kiefer K, et al. Role of type I interferons in the activation of autoreactive B cells. Immunol Cell Biol. 2012;90(5):498–504.

12. Furie R, et al. Anifrolumab, an Anti-Interferon-α Receptor Monoclonal Antibody, in Moderate-to-Severe Systemic Lupus Erythematosus. Arthritis Rheumatol. 2017;69(2):376–86.

13. Houssiau FA, et al. IFN-α kinoid in systemic lupus erythematosus: results from a phase IIb, randomised, placebo-controlled study. Ann Rheum Dis. 2020;79(3):347–55.

14. Pellerin A, et al. Anti-BDCA2 monoclonal antibody inhibits plasmacytoid dendritic cell activation through Fc-dependent and Fc-independent mechanisms. EMBO Mol Med. 2015;7(4):464–76.

15. Goldblatt F, et al. Serious infections in British patients with systemic lupus erythematosus: hospitalisations and mortality. Lupus. 2009;18(8):682–9.

16. Edwards CJ, et al. Hospitalization of individuals with systemic lupus erythematosus: characteristics and predictors of outcome. Lupus. 2003;12(9):672–6.

17. Lee SJ, Silverman E, and Bargman JM. The role of antimalarial agents in the treatment of SLE and lupus nephritis. Nat Rev Nephrol. 2011;7(12):718–29.

18. Kan H, et al. Longitudinal Treatment Patterns and Associated Outcomes in Patients With Newly Diagnosed Systemic Lupus Erythematosus. Clin Ther. 2016;38(3):610–24.

19. Kuznik A, et al. Mechanism of endosomal TLR inhibition by antimalarial drugs and imidazoquinolines. J Immunol. 2011;186(8):4794–804.

20. Häcker H, et al. CpG-DNA-specific activation of antigen-presenting cells requires stress kinase activity and is preceded by non-specific endocytosis and endosomal maturation. EMBO J. 1998;17(21):6230–40.

21. Lau CM, et al. RNA-associated autoantigens activate B cells by combined B cell antigen receptor/Toll-like receptor 7 engagement. J Exp Med. 2005;202(9):1171–7.

22. Seo MR, et al. Hydroxychloroquine treatment during pregnancy in lupus patients is associated with lower risk of preeclampsia. Lupus. 2019;28(6):722–30.

23. Clowse ME, et al. Hydroxychloroquine in lupus pregnancy. Arthritis Rheum. 2006;54(11):3640–7.

24. Alarcón GS, et al. Effect of hydroxychloroquine on the survival of patients with systemic lupus erythematosus: data from LUMINA, a multiethnic US cohort (LUMINA L). Ann Rheum Dis. 2007;66(9):1168–72.

25. Fanouriakis A, et al. 2019 update of the EULAR recommendations for the management of systemic lupus erythematosus. Ann Rheum Dis. 2019;78(6):736−45.

26. Tett SE, et al. Bioavailability of hydroxychloroquine tablets in healthy volunteers. Br J Clin Pharmacol. 1989;27(6):771–9.

27. Liu LH, et al. Understanding Nonadherence with Hydroxychloroquine Therapy in Systemic Lupus Erythematosus. J Rheumatol. 2019;46(10):1309–15.

28. Melles RB, and Marmor MF. The risk of toxic retinopathy in patients on long-term hydroxychloroquine therapy. JAMA Ophthalmol. 2014;132(12):1453–60.

29. Kim JW, et al. Risk of Retinal Toxicity in Longterm Users of Hydroxychloroquine. J Rheumatol. 2017;44(11):1674–9.

30. Rehor A, Hubbell JA, and Tirelli N. Oxidation-sensitive polymeric nanoparticles. Langmuir. 2005;21(1):411–7.

31. Allen SD, et al. Polymersomes scalably fabricated via flash nanoprecipitation are non-toxic in non-human primates and associate with leukocytes in the spleen and kidney following intravenous administration. Nano Research. 2018;11(10):5689–703.

32. Yi S, et al. Tailoring Nanostructure Morphology for Enhanced Targeting of Dendritic Cells in Atherosclerosis. ACS Nano. 2016;10(12):11290–303.

33. Dowling DJ, et al. Toll-like receptor 8 agonist nanoparticles mimic immunomodulating effects of the live BCG vaccine and enhance neonatal innate and adaptive immune responses. J Allergy Clin Immunol. 2017;140(5):1339–50.

34. Geng Y, et al. Shape effects of filaments versus spherical particles in flow and drug delivery. Nat Nanotechnol. 2007;2(4):249–55.

35. Allen S, Vincent M, and Scott E. Rapid, Scalable Assembly and Loading of Bioactive Proteins and Immunostimulants into Diverse Synthetic Nanocarriers Via Flash Nanoprecipitation. J Vis Exp. 2018(138).

36. Allen S, et al. Facile assembly and loading of theranostic polymersomes via multi-impingement flash nanoprecipitation. J Control Release. 2017;262:91–103.

37. Borchert H, et al. Determination of nanocrystal sizes: a comparison of TEM, SAXS, and XRD studies of highly monodisperse CoPt3 particles. Langmuir. 2005;21(5):1931–6.

38. Blanco E, Shen H, and Ferrari M. Principles of nanoparticle design for overcoming biological barriers to drug delivery. Nat Biotechnol. 2015;33(9):941–51.

39. Alexis F, et al. Factors affecting the clearance and biodistribution of polymeric nanoparticles. Mol Pharm. 2008;5(4):505–15.

40. Xiao K, et al. The effect of surface charge on in vivo biodistribution of PEG-oligocholic acid based micellar nanoparticles. Biomaterials. 2011;32(13):3435–46.

41. Patel HM, and Moghimi SM. Serum-mediated recognition of liposomes by phagocytic cells of the reticuloendothelial system - The concept of tissue specificity. Adv Drug Deliv Rev. 1998;32(1-2):45–60.

42. Champion JA, and Mitragotri S. Role of target geometry in phagocytosis. Proc Natl Acad Sci U S A. 2006;103(13):4930–4.

43. Jang CH, et al. Chloroquine inhibits production of TNF-alpha, IL-1beta and IL-6 from lipopolysaccharide-stimulated human monocytes/macrophages by different modes. Rheumatology (Oxford). 2006;45(6):703–10.

44. Karres I, et al. Chloroquine inhibits proinflammatory cytokine release into human whole blood. Am J Physiol. 1998;274(4):R1058–64.

45. Martinson JA, et al. Chloroquine modulates HIV-1-induced plasmacytoid dendritic cell alpha interferon: implication for T-cell activation. Antimicrob Agents Chemother. 2010;54(2):871–81.

46. Bennett L, et al. Interferon and granulopoiesis signatures in systemic lupus erythematosus blood. J Exp Med. 2003;197(6):711–23.

47. Wang X, and Xia Y. Anti-double Stranded DNA Antibodies: Origin, Pathogenicity, and Targeted Therapies. Front Immunol. 2019;10:1667.

48. Tian J, et al. Toll-like receptor 9-dependent activation by DNA-containing immune complexes is mediated by HMGB1 and RAGE. Nat Immunol. 2007;8(5):487–96.

49. Weckerle CE, et al. Network analysis of associations between serum interferon-α activity, autoantibodies, and clinical features in systemic lupus erythematosus. Arthritis Rheum. 2011;63(4):1044–53.

50. Vollmer J, et al. Immune stimulation mediated by autoantigen binding sites within small nuclear RNAs involves Toll-like receptors 7 and 8. J Exp Med. 2005;202(11):1575–85.

51. Gibbert K, et al. IFN-α subtypes: distinct biological activities in anti-viral therapy. Br J Pharmacol. 2013;168(5):1048–58.

52. Rovin BH, et al. Efficacy and safety of voclosporin versus placebo for lupus nephritis (AURORA 1): a double-blind, randomised, multicentre, placebo-controlled, phase 3 trial. Lancet. 2021;397(10289):2070–80.

53. Souyris M, et al. TLR7 escapes X chromosome inactivation in immune cells. Sci Immunol. 2018;3(19).

54. Lande R, et al. Neutrophils activate plasmacytoid dendritic cells by releasing self-DNA-peptide complexes in systemic lupus erythematosus. Sci Transl Med. 2011;3(73):73ra19.

55. Muñoz LE, et al. The role of defective clearance of apoptotic cells in systemic autoimmunity. Nat Rev Rheumatol. 2010;6(5):280–9.

56. Wu YW, Tang W, and Zuo JP. Toll-like receptors: potential targets for lupus treatment. Acta Pharmacol Sin. 2015;36(12):1395–407.

57. Liu Y-J. IPC: professional type 1 interferon-producing cells and plasmacytoid dendritic cell precursors. Annu Rev Immunol. 2005;23:275–306.

58. Tucci M, et al. Glomerular accumulation of plasmacytoid dendritic cells in active lupus nephritis: role of interleukin-18. Arthritis Rheum. 2008;58(1):251–62.

59. Chen K, et al. Tissue-resident dendritic cells and diseases involving dendritic cell malfunction. Int Immunopharmacol. 2016;34:1–15.

60. Kim JH, et al. Intravenously administered gold nanoparticles pass through the blood-retinal barrier depending on the particle size, and induce no retinal toxicity. Nanotechnology. 2009;20(50):505101.

61. Zhu S, et al. Safety Assessment of Nanomaterials to Eyes: An Important but Neglected Issue. Adv Sci (Weinh). 2019;6(16):1802289.

62. Gao H, Shi W, and Freund LB. Mechanics of receptor-mediated endocytosis. Proc Natl Acad Sci U S A. 2005;102(27):9469–74.

63. Swiecki M, and Colonna M. The multifaceted biology of plasmacytoid dendritic cells. Nat Rev Immunol. 2015;15(8):471–85.

64. Zhou D, Kang KH, and Spector SA. Production of interferon α by human immunodeficiency virus type 1 in human plasmacytoid dendritic cells is dependent on induction of autophagy. J Infect Dis. 2012;205(8):1258–67.

65. Al-Hourani K, et al. Innate triggering and antiviral effector functions of activin A. bioRxiv. 2021:2021.03.23.436626.

66. Heil F, et al. Species-specific recognition of single-stranded RNA via toll-like receptor 7 and 8. Science. 2004;303(5663):1526–9.

67. Look M, et al. The nanomaterial-dependent modulation of dendritic cells and its potential influence on therapeutic immunosuppression in lupus. Biomaterials. 2014;35(3):1089–95.

68. Look M, et al. Nanogel-based delivery of mycophenolic acid ameliorates systemic lupus erythematosus in mice. J Clin Invest. 2013;123(4):1741–9.

69. Ganugula R, et al. A highly potent lymphatic system-targeting nanoparticle cyclosporine prevents glomerulonephritis in mouse model of lupus. Sci Adv. 2020;6(24):eabb3900.

70. Hu J, et al. A novel long-acting azathioprine polyhydroxyalkanoate nanoparticle enhances treatment efficacy for systemic lupus erythematosus with reduced side effects. Nanoscale. 2020;12(19):10799–808.

71. Yuan F, et al. Dexamethasone prodrug treatment prevents nephritis in lupus-prone (NZB × NZW)F1 mice without causing systemic side effects. Arthritis Rheum. 2012;64(12):4029–39.

72. Bennett L, et al. Interferon and granulopoiesis signatures in systemic lupus erythematosus blood. J Exp Med. 2003;197(6):711–23.

73. Zhuang H, et al. Animal Models of Interferon Signature Positive Lupus. Front Immunol. 2015;6:291.

74. Reeves WH, et al. Induction of autoimmunity by pristane and other naturally occurring hydrocarbons. Trends Immunol. 2009;30(9):455–64.

75. Bobbala S, Allen SD, and Scott EA. Flash nanoprecipitation permits versatile assembly and loading of polymeric bicontinuous cubic nanospheres. Nanoscale. 2018;10(11):5078–88.

76. Karabin NB, et al. The Combination of Morphology and Surface Chemistry Defines the Biological Identity of Nanocarriers in Human Blood. bioRxiv. 2020:2020.09.02.280404.

77. Cerritelli S, et al. Aggregation behavior of poly(ethylene glycol-bl-propylene sulfide) di- and triblock copolymers in aqueous solution. Langmuir. 2009;25(19):11328–35.

78. McCall RL, and Sirianni RW. PLGA nanoparticles formed by single- or double-emulsion with vitamin E-TPGS. J Vis Exp. 2013(82):51015.

79. Karabin NB, et al. Sustained micellar delivery via inducible transitions in nanostructure morphology. Nat Commun. 2018;9(1):624.

80. Krug A, et al. Identification of CpG oligonucleotide sequences with high induction of IFN-alpha/beta in plasmacytoid dendritic cells. Eur J Immunol. 2001;31(7):2154–63.

