## Supplemental Data for "Targeted Delivery of Chloroquine to Plasmacytoid Dendritic Cells Enhances Inhibition of the Type I Interferon Response"

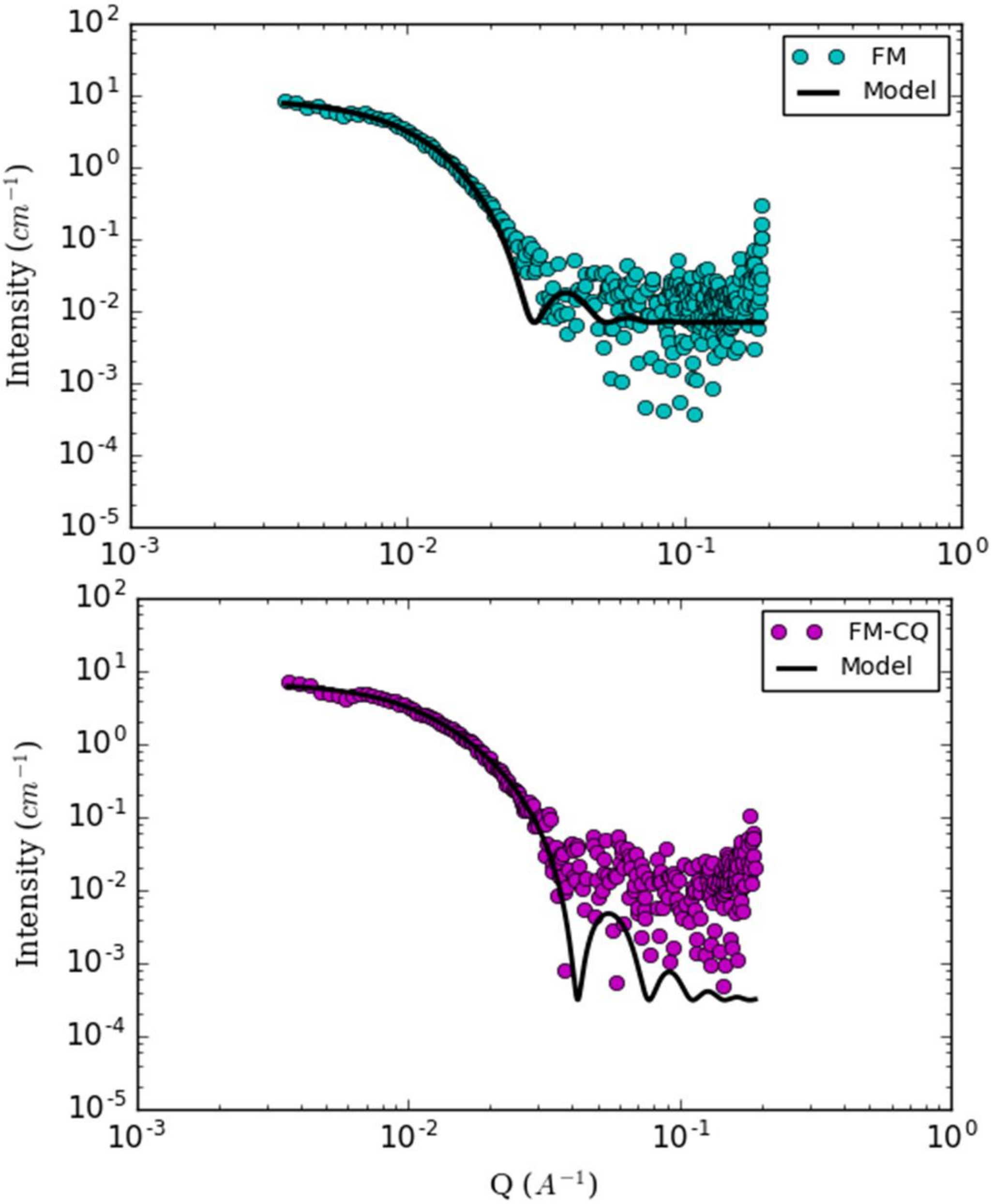


**Figure S1. SAXS model fit for blank and CQ-loaded FMs.** Unloaded ($\chi^2$ = 0.008) and CQ-loaded FM ($\chi^2$ = 0.0012) characteristics were determined by fitting the scattering profiles with a flexible cylindrical model using SASVIEW 4.X. Model analysis used the following parameters: 2 µm cylinder length, 150 nm persistence length, and 8 nm PPS core radius.

A

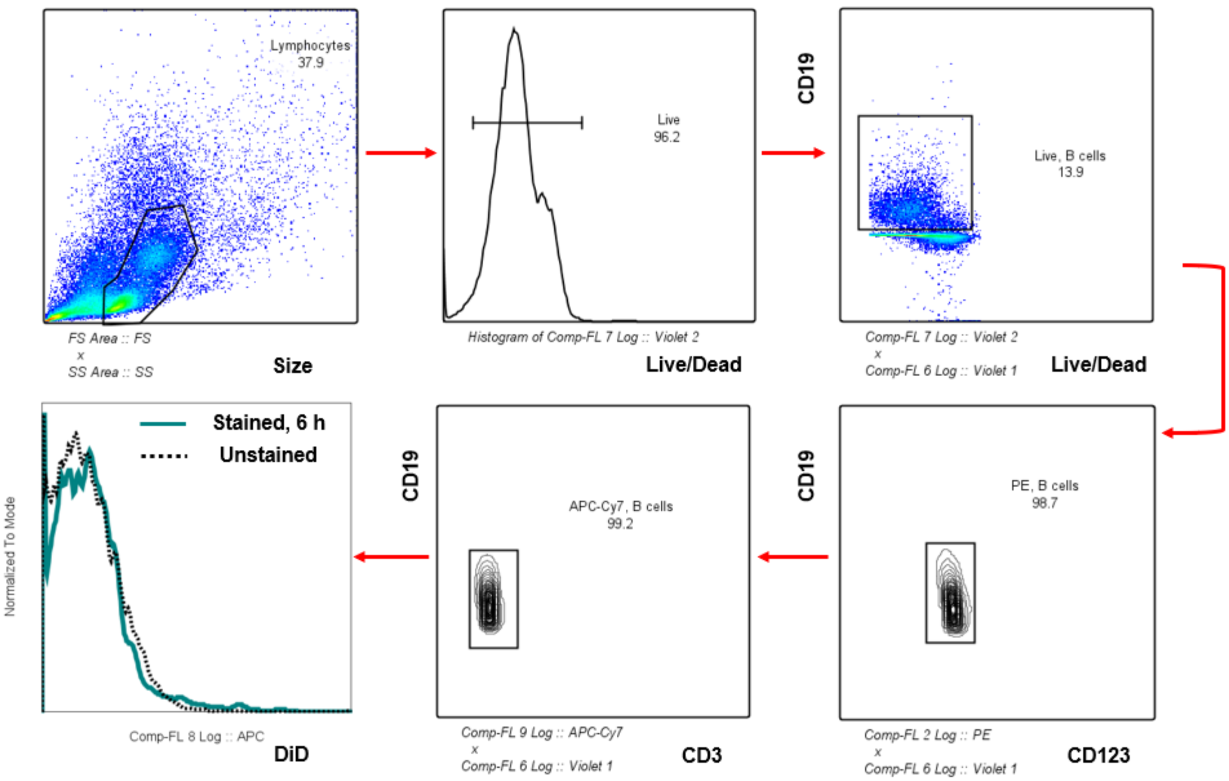

Lymphocytes
37.9
FS Area :: FS
x
SS Area :: SS
Size
Live
96.2
Histogram of Comp-FL 7 Log :: Violet 2
Live/Dead
CD19
Live, B cells
13.9
Comp-FL 7 Log :: Violet 2
x
Comp-FL 6 Log :: Violet 1
Live/Dead
Stained, 6 h
Unstained
Normalized To Mode
Comp-FL 8 Log :: APC
DiD
CD19
APC-Cy7, B cells
99.2
Comp-FL 9 Log :: APC-Cy7
x
Comp-FL 6 Log :: Violet 1
CD3
CD19
PE, B cells
98.7
Comp-FL 2 Log :: PE
x
Comp-FL 6 Log :: Violet 1
CD123


B

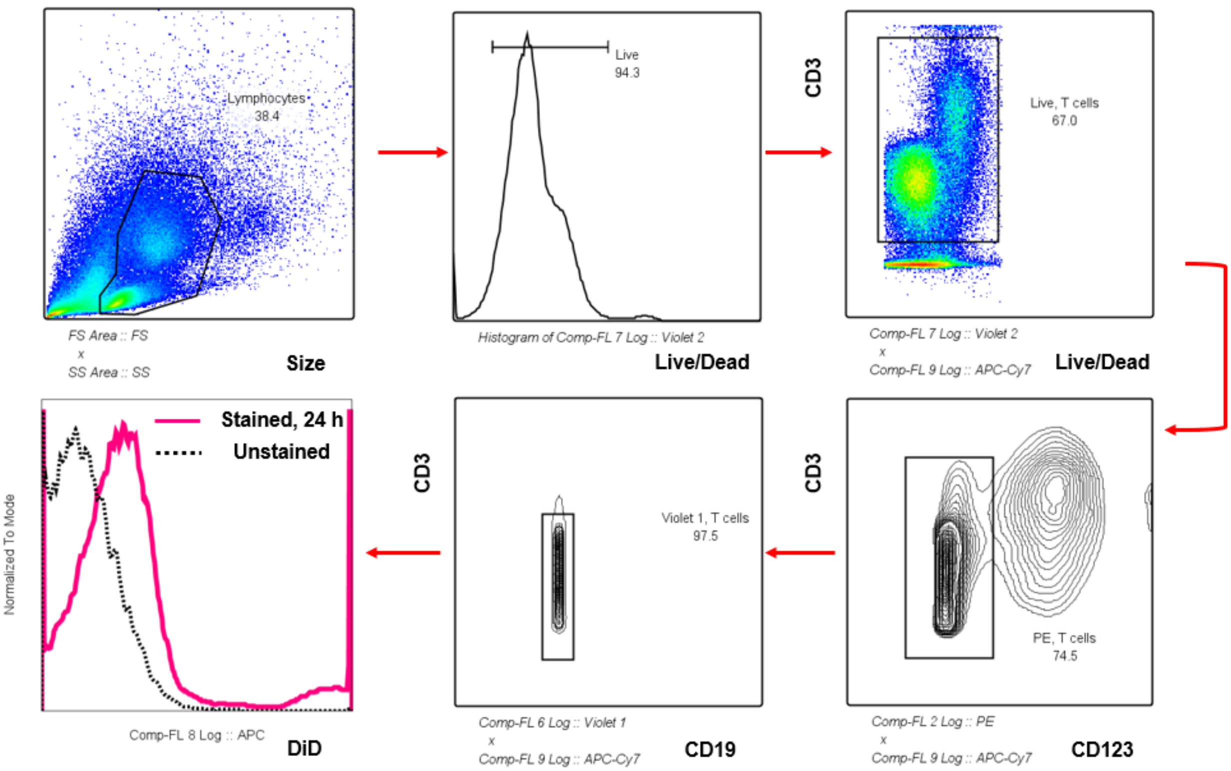

Lymphocytes
38.4
FS Area :: FS
x
SS Area :: SS
Size
Live
94.3
Histogram of Comp-FL 7 Log :: Violet 2
Live/Dead
CD3
Live, T cells
67.0
Comp-FL 7 Log :: Violet 2
x
Comp-FL 9 Log :: APC-Cy7
Live/Dead
Stained, 24 h
Unstained
Normalized To Mode
Comp-FL 8 Log :: APC
DiD
CD3
Violet 1, T cells
97.5
Comp-FL 6 Log :: Violet 1
x
Comp-FL 9 Log :: APC-Cy7
CD19
CD3
PE, T cells
74.5
Comp-FL 2 Log :: PE
x
Comp-FL 9 Log :: APC-Cy7
CD123

C

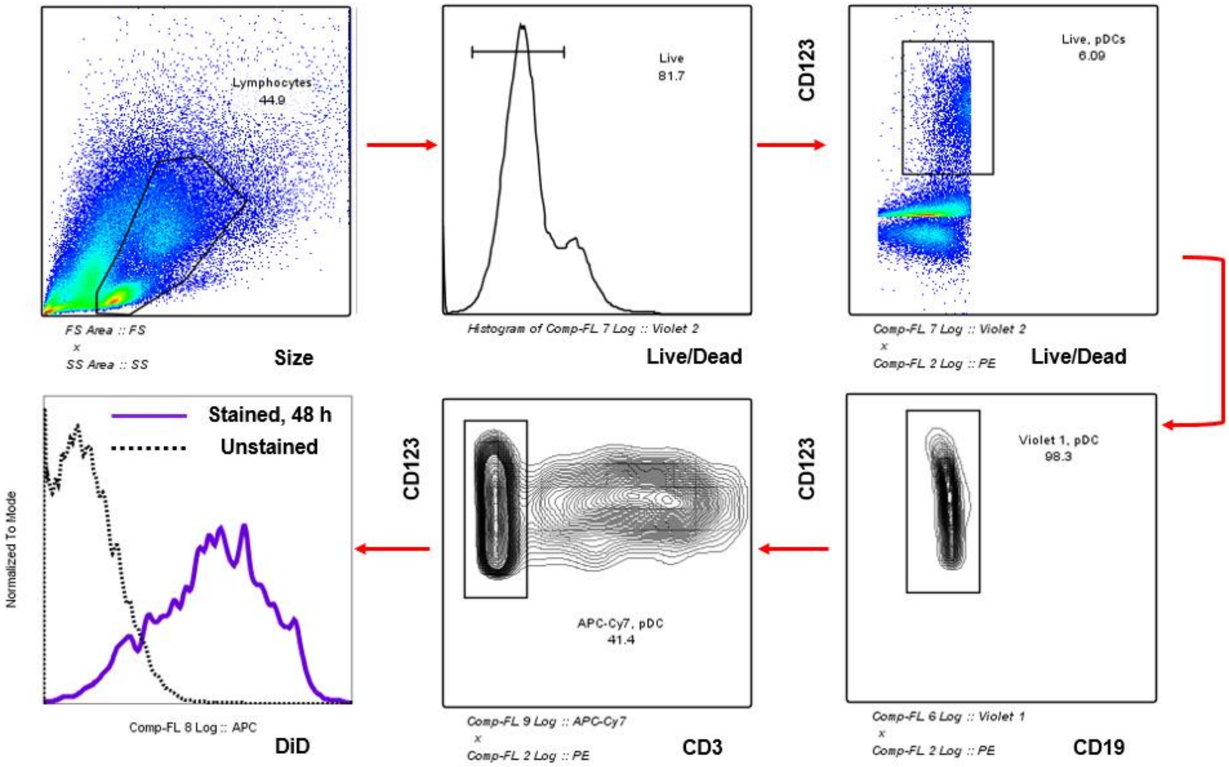


**Figure S2. Representative flow cytometry gating strategy for analysis of DiD-loaded FMs cultured with human PBMCs.** Fresh human PBMCs cultured with 200 µg/mL DiD-loaded FMs in supplemented RPMI GlutaMAX medium plus 10% FBS and 20 ng/mL recombinant human IL-3 were stained with phenotypic cell surface markers: A) CD19+ B cells, B) CD3+ T cells, and C) CD123+ pDCs. DiD+ immune cells were determined as above and independent of other cell surface markers. Median fluorescence intensity of DiD was calculated for each cell type.

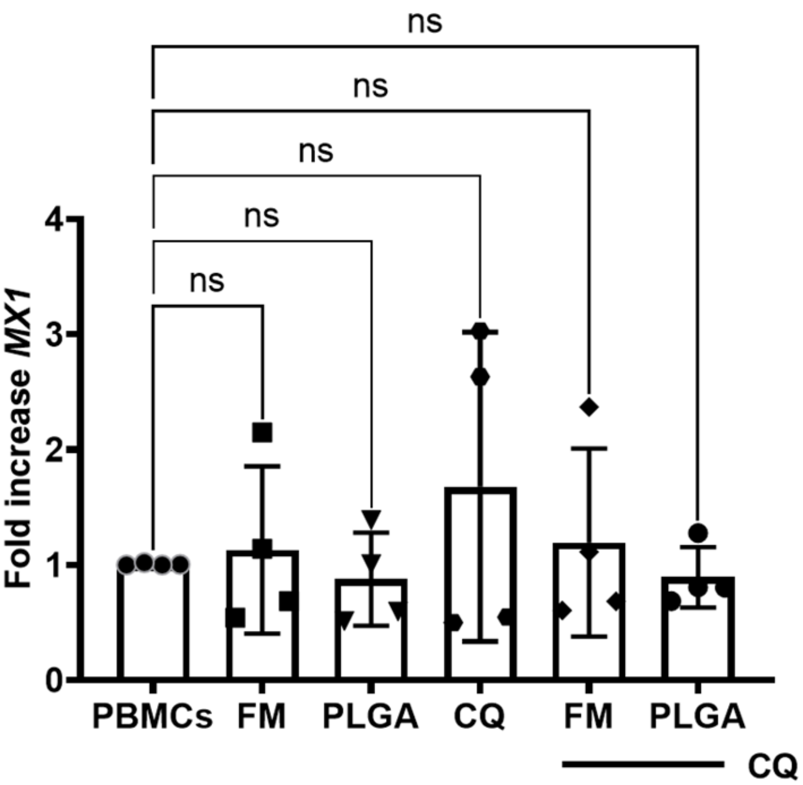


**Figure S3. No significant difference between *MX1* gene expression of empty nanocarriers, CQ alone, and empty nanocarriers with soluble CQ in human PBMCs.** Fresh, healthy human PBMCs were isolated and treated with empty FMs or PLGA nanocarriers with or without soluble CQ at 3.91 μM for 1 h. Total RNA was isolated and *MX1* expression quantified using TaqMan real-time RT-qPCR normalized to β-actin expression. Data are means ± standard deviation for *n* = 4 independent donors. Statistical analysis: One-way ANOVA with Tukey’s multiple comparisons test.
